# Enhancing salt tolerance in a hybrid poplar *(Populus nigra x maximowiczii)* through exogenous putrescine application: Growth, physiological, and biochemical responses

**DOI:** 10.1101/2025.06.04.657897

**Authors:** Sanchari Kundu, Medini Weerasinghe, Maegan Gagne, Subhash Minocha

**Affiliations:** Department of Biological Science, University of New Hampshire, Durham, NH, USA; Present address: School of Plant and Environmental Sciences, Virginia Tech, Blacksburg, VA, USA

**Keywords:** Polyamines, putrescine, foliar spray, salt stress, hybrid poplar, metabolism, growth, amino acids

## Abstract

**Introduction:** Putrescine, a polyamine involved in plant growth and stress responses, has shown potential in mitigating abiotic stress effects. However, little is known about the exogenous addition of putrescine regarding salt tolerance in trees.

**Methods:** This study was conducted to investigate whether exogenous putrescine application via foliar spray enhances growth in a hybrid poplar (*Populus nigra x maximowiczii,* clone NM6) under a short duration of salt stress. Salt stress was induced by irrigating roots with 100 mM and 200 mM NaCl, followed by foliar spraying of putrescine on several days. Measurement of growth including plant height and stem diameter for each plant were recorded in the greenhouse every 15 days throughout the experiment. Gas exchange, total chlorophyll, carotenoids, soluble sugars and proteins, amino acids, polyamines, and relative water content were analyzed in foliage collected 3, 6, 7, 13, 20, 35 days after treatment.

**Results:** Putrescine spray on the salt-treated plants caused a significant increase in plant growth. Putrescine application caused a significant increase in fructose, glucose, and galactose in all plants, but putrescine spray had a variable impact on the sucrose content of 100 mM NaCl-treated plants. Based on metabolic responses, plants treated with 100 mM NaCl fared better when sprayed with putrescine than those treated with 200 mM NaCl.

**Discussion:** Exogenous application of putrescine mitigated growth inhibition effects of salinity. These finding highlight the potential of putrescine as a practical approach to increase salt tolerance in young poplar trees, with implications for forestry and land reclamation in saline environment.

## 1 Introduction

Polyamines (PAs), such as putrescine (Put), spermidine (Spd), and spermine (Spm), are small aliphatic amines derived from amino acids (AAs) that play essential roles in plant development and adaptation to abiotic stress (Alcázar et al., 2006, Minocha et al., 2014, Pál et al., 2018, Atabayeva et al., 2020, Antoniou et al., 2021, Shao et al., 2022, Borromeo et al., 2023, Blázquez, 2024, Tabur et al., 2024, Zhao et al., 2024b). They influence diverse physiological processes such as seed germination, photosynthesis, root architecture, and cellular signaling. In recent years, PAs have attracted attention for their involvement in enhancing stress tolerance through metabolic and physiological modulation (Wuddineh et al., 2017, Antoniou et al., 2021, Imran et al., 2021, Sundararajan et al., 2021, Borromeo et al., 2023, Pál et al., 2025). However, their role in woody plants-particularly fast-growing poplars, remains unexplored despite their significant ecological and economic importance (Zalesny Jr et al., 2019). Long life cycles, complex anatomical structures, and ethical considerations in genetic studies, all create unique challenges in research with woody plants (Neale and Kremer, 2011, Chen et al., 2021). Understanding the stress adaptation mechanisms in woody perennial plants is crucial for sustainable forestry and climate resilience, particularly in response to global warming.

Among various abiotic stresses, salinity is a major factor that disrupts plant growth, photosynthesis, ion homeostasis, water balance, carbon and nitrogen metabolism (Bargmann et al., 2009, Cramer et al., 2011, Dinneny, 2015, Parihar et al., 2015, Hanson et al., 2016, Haak et al., 2017, Zhang et al., 2020a, Sun et al., 2022, Calhoun et al., 2023, Kundu, 2023, Zhang et al., 2023). Azimi et al. (2021) reported that N deficiency is a major consequence of salt stress, reducing leaf area and inducing chlorosis, which hinders plant productivity. While N fertilization is a common approach for mitigation, run-off from excessive application of N can contribute to environmental issues such as water pollution and algal blooms, necessitating more feasible stress mitigation strategies (Li et al., 2021, Zhang et al., 2020a).

Polyamines have been recognized as key regulators of abiotic stress responses, accumulating in plants subjected to salinity, heavy metals, drought, and temperature extremes (Alcázar et al., 2010, Mohapatra et al., 2010, Pál et al., 2018, Malik et al., 2022, Kundu, 2023, Wang et al., 2024b, Liu et al., 2025b). They play a vital role in modulating carbon metabolism, stabilizing membrane integrity, scavenging reactive oxygen species (ROS), and regulating osmotic balance (Majumdar et al., 2013, Pál et al., 2021, Pál et al., 2025). Polyamines can also enhance N assimilation, thereby improving chlorophyll synthesis and overall plant growth (Alcázar et al., 2010, Tamang et al., 2021, Zhang et al., 2024). Several studies have demonstrated that exogenous PA application improves plant resilience by modulating hormone signaling like abscisic acid, gene expression, and enzymatic responses under salt stress and others (Paul et al., 2018, Xiong et al., 2018, Borromeo et al., 2023, Wang et al., 2024b). Seed priming with PAs has been reported to enhance photosynthetic pigments, proline accumulation, root growth and biomass production in crops such as soybean, rapeseed, tomato, and rice (Sheteiwy et al., 2021, ElSayed et al., 2022, Borromeo et al., 2023, Zhang et al., 2024) under salinity stress.

Additionally, exogenous PA treatments have been shown to increase soluble sugars, proteins, and amino acid accumulation, enhancing abiotic stress tolerance in economically significant plants like grapevine, ginseng, rice, and tomato (Sundararajan et al., 2021). Recent studies have also shown that exogenous Spd treatment increased resistance to fusariosis in flax by suppression of PA metabolism (Augustyniak et al., 2025). However, despite these promising findings, studies in woody perennials remain limited, particularly in fast-growing trees like poplars.

Poplars (*Populus* spp.) are among the fastest-growing perennial tree species, widely distributed across North America, China and India (Dickmann, 2001, Hawkins et al., 2003, Plomion et al., 2016, Zalesny Jr et al., 2019, Buell et al., 2023); and are key resources used in sustainable forestry, bioenergy production, and the lumber industry (Yi et al., 2022). Poplars exhibit notable stress tolerance, including resistance to salt and heavy metal contamination (Zalesny Jr et al., 2019, Lin et al., 2023), making them ideal candidates for genetic and physiological research (Guerra et al., 2011). Several varieties of poplars have shown potential to be grown in marginal sandy lands (Ghezehei et al., 2021). Advances in’omics’ technologies and the sequencing of the poplar genome (Ma et al., 2019) have enabled deeper insights into their stress response pathways. Notably, transgenic studies in hybrid poplars have identified genes like *NAC13,* which enable salt tolerance (Zhang et al., 2019). Some recent studies have applied single-cell RNA-seq transcriptomics (scRNA-seq) to dissect poplar vascular root system and adaptations in response to soil compaction and abiotic stress, revealing cell-type-specific regulatory mechanisms (Conde et al., 2022, Liu et al., Li et al., 2024, Zhu et al., 2025). Recent computational advances have further enabled cross-species comparison of conserved cell types, expanding the potential for identifying PA-mediated functions across plant taxa at single-cell resolution (Chau et al., 2025). Although not included in this study, such approaches offer promising avenues for future research on PA specific stress responses in woody perennials. Understanding how PAs contribute to salt stress tolerance at tissue-or cell-specific resolution could bridge a critical knowledge gap in tree physiology and enhance strategies for improving stress resilience in forest trees.

Among poplar hybrids, *Populus nigra x maximowiczii* (NM6 clone) is widely cultivated in North America due to its fast growth, high biomass yield, asexual propagation capacity, and genetic stability (Labrecque and Teodorescu, 2005, Guerra et al., 2011). Zalesny Jr et al. (2019) also reported that the NM6 clone was more salt-tolerant than other poplar varieties. Yet no studies have investigated the physiological and biochemical effects of exogenous PA application in mitigating salt stress in NM6 poplars. Given the increasing soil salinity issues affecting forestry and agricultural productivity, understanding how PA treatments effects stress tolerance in NM6 poplars is of scientific as well as economic significance.

To address this research gap, we investigated the role of exogenous PA application: specifically Put in enhancing salt stress tolerance in young NM6 poplars. In a controlled greenhouse experiment, we assessed the physiological and biochemical responses of NM6 poplar cuttings to NaCl-induced stress and examined whether foliar-application of Put could alleviate its adverse effects. The general objective of this study was to determine how exogenous application of Put influences plant responses to salt stress, with a focus on physiological traits, photosynthetic performance, and key metabolic indicators. To guide this investigation, we formulated the following research questions: 1. How does NaCl stress affect the physiological and metabolic processes in NM6 poplar plants? 2. Can the exogenous application of Put mitigate NaCl-induced stress; and if so, through which biochemical and physiological pathways? We hypothesized that foliar application of Put would enhance salt stress tolerance in NM6 poplars by improving plant growth, photosynthetic efficiency, and metabolic stability (soluble sugars, AAs, and PAs) thereby promoting growth and stress resilience. These improvements are expected to promote overall plant resilience under saline conditions. This study contributes novel insights into PA-mediated stress mitigation in woody plants and advances sustainable strategies to improve abiotic stress in forestry species.

## 2 Materials and methods

### 2.1 Plant Material and Growth Conditions

Cuttings of hybrid poplar (*Populus nigra x maximowiczii* - NM6) plant were collected from a 5-year-old, healthy tree at the University of New Hampshire (UNH) Kingman Farm in Durham (43.1725143,-70.9462316). The cuttings were rooted and maintained at the UNH MacFarlane greenhouse from mid-April to late June 2021 as an acclimatization period prior to the start of experimental treatments in early July.

During the initial growth phase, cuttings (∼15 cm in height, 0.6 cm stem diameter) were kept under mist for two weeks in 17.78 cm grow tubes containing PRO-MIX® soil with Mycorrhizae peat based growing medium (Premier Tech, Quebec, Cananda). Each 17.78 cm grow tube (approximately 5 cm in diameter) was filled with an estimated 349 cm³ of PRO-MIX® soil with Mycorrhizae. The grow tubes were placed in trays to retain excess water, ensuring consistent soil moisture, and plants were watered regularly. After 2 months, well-established plants (∼30–38 cm in height with 9–10 leaves) were selected and transferred to 33 cm pots filled with a 1:1 mixture of vermiculite and perlite.

The experiments were conducted from July to September 2021 in a controlled greenhouse environment with natural sunlight and a 16-hour photoperiod. The temperature in the greenhouse ranged between 22°C and 24°C, with a relative humidity of 70%. Plants were irrigated and fertilized twice daily using an automated drip-line irrigation system with Jack’s Pure Water LX - Professional fertilizer® (J.R. Peters Inc., Allentown, PA, USA). Each plant received 200 mL of water at 08:00 AM and 2:00 PM, and additional watering with plain water was applied manually in the evening when needed.

### 2.2 Experimental Design

A completely fixed design was used in this study, ensuring that plants remained in fixed positions throughout the experiment. One time salt treatment with sodium chloride (NaCl) was added 21 days after transplanting the cuttings into 33 cm pots, when the plants were approximately 3 months after the initial collection of cuttings. There was a total of 36 cuttings, 6 treatment assigned to 6 treatment groups (n = 6 per treatment): treatment A (control-no NaCl, no Put), treatment B (100 mM NaCl), treatment C (200 mM NaCl), treatment D (control + 1 mM Put foliar spray), treatment E (100 mM NaCl + 1 mM Put foliar spray), and treatment F (200 mM NaCl + 1 mM Put foliar spray).

A 0.5% Silwet^TM^ (Momentive Performance Materials, Niskayuna, NY, USA) solution was added as a surfactant to enhance the penetration of the 1 mM Putrescine (dihydrochloride ≥ 98%; Sigma-Aldrich, Inc., St. Louis, MO, USA) foliar spray.. To ensure uniform salt application, drip irrigation was halted 18 hours before salt treatment. Salt treatments were delivered via root irrigation by hand-pouring 200 mL of NaCl solution per pot at the time zero of treatment (Day 0) and again 6 hours later. Drip irrigation resumed 6 hours after the final salt application. Saucers were placed beneath the pots to prevent water leakage.

Thirty milliliters of 1 mM Put foliar spray was applied to cuttings in treatment groups D, E, and F immediately following the salt treatment. The spray was applied only to the leaves while the soil was carefully covered with plastic sheets to prevent contamination. After spraying, any excess solution was manually tapped off the leaves. Putrescine spray occurred at time 0, and days 3, 6, and 13.

### 2.3 Sample Collection

Plant samples including the remaining leaves and root tissues, were collected at various time points before and after treatment application for physiological and biochemical analyses.

### 2.3a Leaf Sampling

All cuttings were sampled at time zero and one representative plant per treatment group was sampled 3, 6, 13, and 20 days post treatment. For each sampling, the fully expanded 6^th^ leaf from the plant apex was selected, washed with fresh water and patted dry between 2 layers of paper towels. From those leaves ∼6 mm disks were punched using a common paper punch to create a pool of ∼2 g. The leaf disks were mixed, and sub-samples were taken for various analyses; each placed into pre-weighed 2 ml microfuge tubes and appropriate volumes and buffers added. On the day of collection, all samples were collected and transported on ice and later stored at-20°C until further analysis.

Physiological traits like relative water content (RWC) and chlorophyll content were measured using leaf discs. Gas exchange measurements were taken with LICOR-6400 directly on the plant leaves. Biochemical assays like soluble sugars, PAs, AAs, and total protein were done with leaf discs. All samples (except leaf discs for biomass measurements) were stored at-20 °C until further analyses.

### 2.3b Roots Sampling

Root samples were collected from 4 biological replicates per treatment only once at the time of harvesting. Roots were thoroughly washed under running tap water to remove residual vermiculite and perlite and pat-dried. Fine secondary roots (2–3 mm in diameter) were carefully snipped from the main root using sterilized scissors. A pool of ∼50 mg of fresh root tissue was collected, mixed thoroughly, places into a pre-weighed 2 mL microfuge tubes and 1 ml 5% perchloric acid (HCLO_4_) was added. All root samples were collected and transported on ice and later stored at-20°C until further analysis.

### 2.4 Plant Growth Parameters

Plant height and stem diameter measurements were taken from 6 replicates per treatment and recorded at time 0 and on days 15, 30, and 45 post salt treatment. Basal diameters were measured using a digital caliper, with a consistent reference point marked near the base of the stem to ensure accuracy across measurement intervals. Growth in height and stem diameter was expressed as a percentage increase using the following formulas:

Increase in height (%) = [(Height on next day - height on previous day)/ Height of day 1] * 100 Increase in diameter (%) = [Diameter on next day – diameter on previous day)/ Diameter of day 1] * 100

### 2.5 Soluble Sugars

*Sample preparation:* Soluble sugars were quantified following a modified protocol from Blagden et al. (2022). Leaf discs (50 ± 2 mg FW) were incubated at 65°C for 30 min in 1 mL of 80% ethanol.

*Analysis:* A total of 11 sugars were quantified: xylose, arabinose, fructose, mannose, glucose, galactose, sucrose, trehalose, rhamnose monohydrate, maltose monohydrate, and raffinose pentahydrate. Detection was performed using a Shimadzu RID-10A refractive index detector (RID) set at 30°C (Shimadzu Scientific Instruments Inc., Columbia, MD). The total run time was 15 min, including column washing and stabilization between injections. The column temperature was maintained at 25°C throughout the analysis. Chromatographic data were processed using Perkin Elmer TotalChrom software (version 6.2.1). Peaks were identified by matching retention times with known sugar standards. An 8-point external standard curve (3 mg mL ¹) was generated for individual sugar quantification (Supplemental Table 2). Unresolved sugar peaks were quantified as combined concentrations based on their summed peak areas, including xylose + arabinose, glucose + galactose, and trehalose + maltose.

### 2.6 Polyamines and Amino acids

*Sample preparation:* the quantification of different AAs and PAs, approximately 40±2 mg fresh leaf discs were collected in 5% perchloric acid at a ratio of 1:25 (w:v) in 2 mL microfuge tubes. The samples were freeze-thawed three times and processed for dansylation following a modified protocol from Minocha et al. (1994), (Minocha and Long, 2004) and (McDermot et al., 2020).

After the final thawing, the samples were vortexed at high speed for 2 min and centrifuged at 14,000 × g for 8 min. For each sample and external standards, 20 µL of an internal standards mix containing 0.05 mM α-methyl-DL phenylalanine (for AAs) and 0.05 mm heptane diamine (for PAs), both dissolved in 5% HCLO_4_, was added to each tube. Subsequently, 100 µL of the freeze-thawed extract was combined with 100 µL of 2.691 M sodium carbonate and 100 µL of freshly prepared dansyl chloride (20 mg/mL in acetone). The mixture was vortexed and incubated at 60°C for 30 min. After incubation, the microfuge tubes were cooled at room temperature for 3 min, followed by the addition of 45 µL glacial acetic acid to terminate the reaction. Acetone was evaporated using a speed-vac for 10 min, and 1735 µL methanol was added to the mixture. The methanol extract was filtered using a 0.45 µm nylon syringe filter fitted onto a 3 mL syringe before transferring the solution into autosampler vials.

*Analysis:* Separation of AAs and PAs was performed using a 15 cm column (Phenomenex Synergi Hydro-RP 80 Å, LC Column 150 x 4.6 mm, 4 µm). Quantification was achieved using a fluorescence detector (Series 200 PerkinElmer) set at 340 nm for excitation and 510 nm for emission. Relative quantification of AAs and PAs was performed using an external standard curve (Supplemental Table 3). Chromatographic data were analyzed using Perkin Elmer TotalChrom software (version 6.2.1), incorporating a multiplication factor within the software to determine the concentration of each component in nmol g ¹ FW of tissue.

### 2.7 Total Soluble Proteins

Total soluble protein content was quantified following a modified protocol from Minocha et al. (2019). Fresh leaf discs (50 ± 2 mg FW) were extracted in a freshly prepared 100 mM Tris buffer (pH 8.0) containing 20 mM MgCl, 10 mM NaHCO, 1 mM ethylenediaminetetraacetic acid (EDTA), and 10% (v/v) glycerol. The extraction was performed using a three-cycle freeze-thaw method.

The extract was centrifuged at 13,000 × g for 5 min, and the supernatant was used for total soluble protein quantification following the Bradford (1976) method. The analysis was conducted using Bio-Rad protein assay dye reagent (Bio-Rad Laboratories, Hercules, CA), with bovine serum albumin (BSA) as the standard. Absorbance was recorded at 595 nm using a spectrophotometer. A standard curve (0.1–0.5 mg/mL) was generated to quantify protein concentrations in the samples.

### 2.8 Relative Water Content

Relative water content (RWC) was determined by measuring the (FW) of the leaf tissue immediately after collection. For DW, the leaf samples were incubated in an oven at 70°C for 48 hours. The samples were reweighed after 24 hours to confirm that no residual moisture remained. Relative water content was calculated using the following formula:

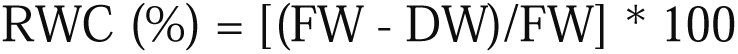

### 2.9 Chlorophyll Content

Chlorophyll content was quantified following the method described in Minocha et al., (2009) and McDermot et al. (2020). Total chlorophyll, chlorophyll a, chlorophyll b, and carotenoids were calculated using equations found in Lichtenthaler (1987).

### 2.10 Leaf Gas Exchange

Gas exchange parameters, including photosynthetic rate (P_n_), transpiration rate (E), and stomatal conductance (g_s_) were measured on the 6^th^ fully expanded leaf from the apex for each cutting.

Measurements were taken on 3–4 plants per treatment using a portable Li-COR photosynthesis system (LI-6400/XT, Li-COR Biosciences, Lincoln, NE, USA).

Gas exchange was recorded between 09:00 and 12:00 under a photosynthetic photon flux density (PPFD) of 1000 μmol m ² s ¹ provided by a red-blue LED light source (6 cm² chamber).

Airflow was maintained at 500 μmol s ¹, and reference CO concentrations were set to 400 μmol mol ¹. The block temperature was maintained at 25°C. Leaf humidity was not actively controlled but ranged between 50–60% during measurements.

### 2.11 Statistical Analysis

All statistical analyses were conducted with JMP® Pro 18 (JMP Statistical Discovery LLC, Cary, NC, USA).

Data were analyzed using a two-way analysis of variance (ANOVA) to assess statistically significant differences between treatments within each time point. All results are presented using the original, untransformed data. Time and treatment were treated as fixed effects. Post-hoc comparisons were performed using Tukey’s test, with statistical significance set at p < 0.05.

Graphical figures of the data were generated using the ggplot2 package, and both heatmap and correlation plots were produced with the pheatmap and corrplot packages in RStudio (R version 4.4.1; Posit, Boston, MA, USA).

## 3 Results

### 3.1 Effects of Put spray on Plant Growth and Morphological Symptoms

Plant height and stem diameter were measured 15, 30 and 45 days after salt treatment. Figure 1 (B, C) shows that both stem length and diameter in the Control and 100 mM NaCl treated cuttings sprayed with foliar Put are greater than either of those treatments without Put spray.

**Figure 1:**
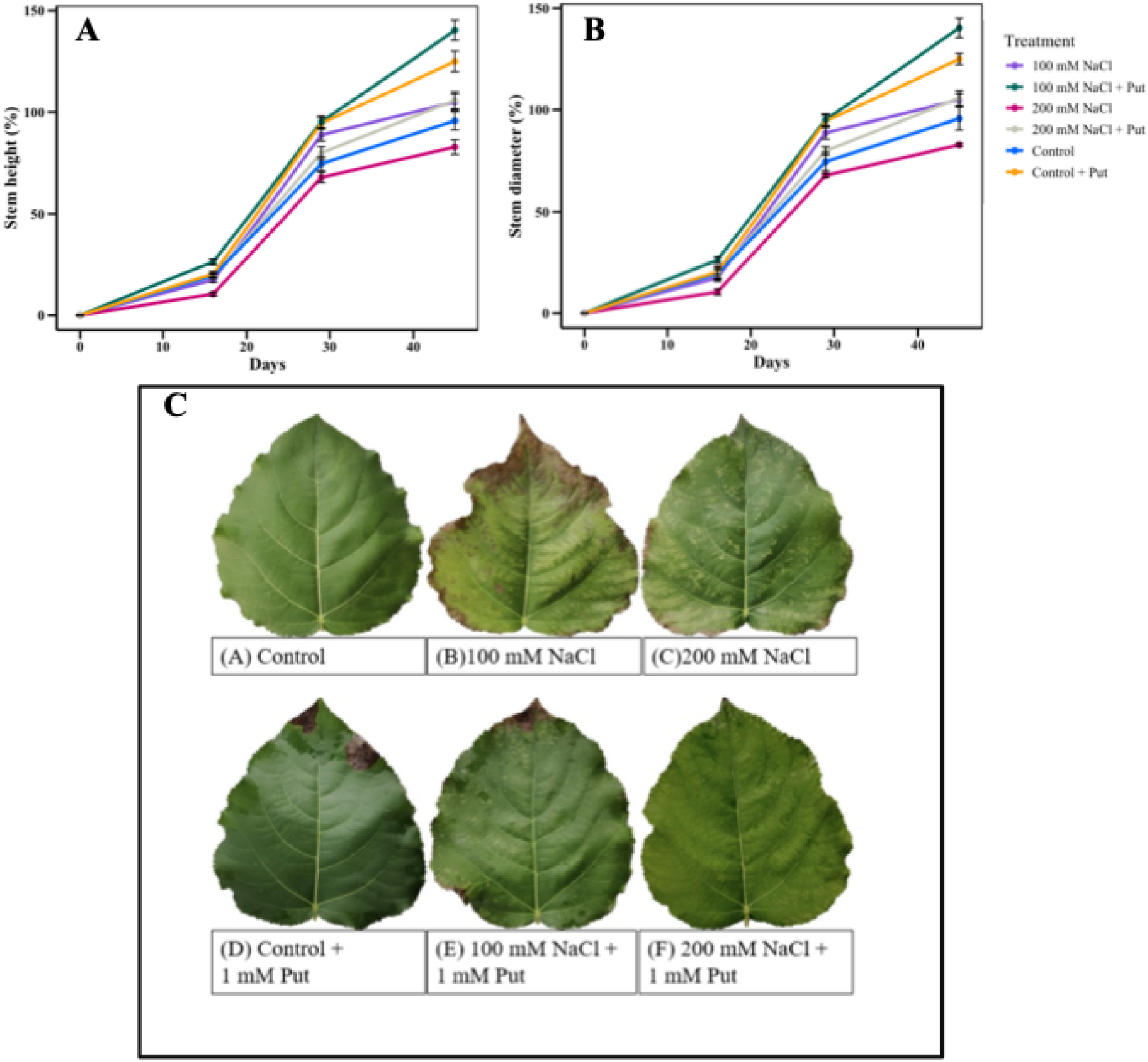
The effect of two different concentrations of NaCl (± Putrescine spray) on the growth and morphology of hybrid poplar NM6 plants. (A, B) Average stem length and stem diameter changes over 45 days of salt treatment. (C) Morphological symptoms of leaves under different treatments: (A) Control, (B) 100 mM NaCl, (C) 200 mM NaCl, (D) Control + 1 mM Put, (E) 100 mM NaCl + 1 mM Put, (F) 200 mM NaCl + 1 mM Put. Data represent mean ± SE (n = 6).

That suggests that under control or low salt conditions, Put had a growth-promoting effect on the poplar cuttings. Foliar discoloration (Fig 1A) began 13 days after salt exposure; leaves of the 100 mM and 200 mM NaCl treated cuttings appear to have a greater degree of chlorosis and necrosis as compared to the control. The application of 1 mM Put reduced these stress symptoms in the salt-treated plants, and all Putrescine-treated leaves displayed a healthier appearance, with markedly less chlorosis and/or necrosis.

### 3.2 Effects of Put spray on Gas exchange, chlorophyll and total soluble protein

Gas exchange parameters, carotenoid levels, and protein content were measured in hybrid poplar NM6 leaves across multiple time points after NaCl and Put treatment. Stomatal conductance (gs) was significantly affected by treatment on day 35. (Fig 2A). On day 35, gs in the Put-treated 100 mM NaCl plants reached 0.55± 0.03 mol m ² s ¹, which was significantly higher than in the control (0.230 ± 0.053 mol m ² s ¹; p < 0.05). The 200 mM NaCl + Put group also showed elevated gs compared to NaCl alone, though not statistically significant (p > 0.05). On day 7, no significant differences were observed among treatments. Transpiration rate (*E*) showed similar trends (Fig. 2D). On day 35, plants treated with 100 mM NaCl + Put had significantly higher *E* (∼6.29 ± 0.2 mmol m ² s ¹) compared to the control (∼5.1 ± 0.3 mmol m ² s ¹) and 100 mM NaClalone (∼3.79 ± 0.61 mmol m ² s ¹; *p* < 0.05). Net photosynthesis (*Pn*), while not statistically different (see Supplementary Fig. 1C), consistently trended 62% higher in Put-treated plants under both 100 mM and 200 mM NaCl compared to untreated salt-stressed groups.

**Figure 2.**
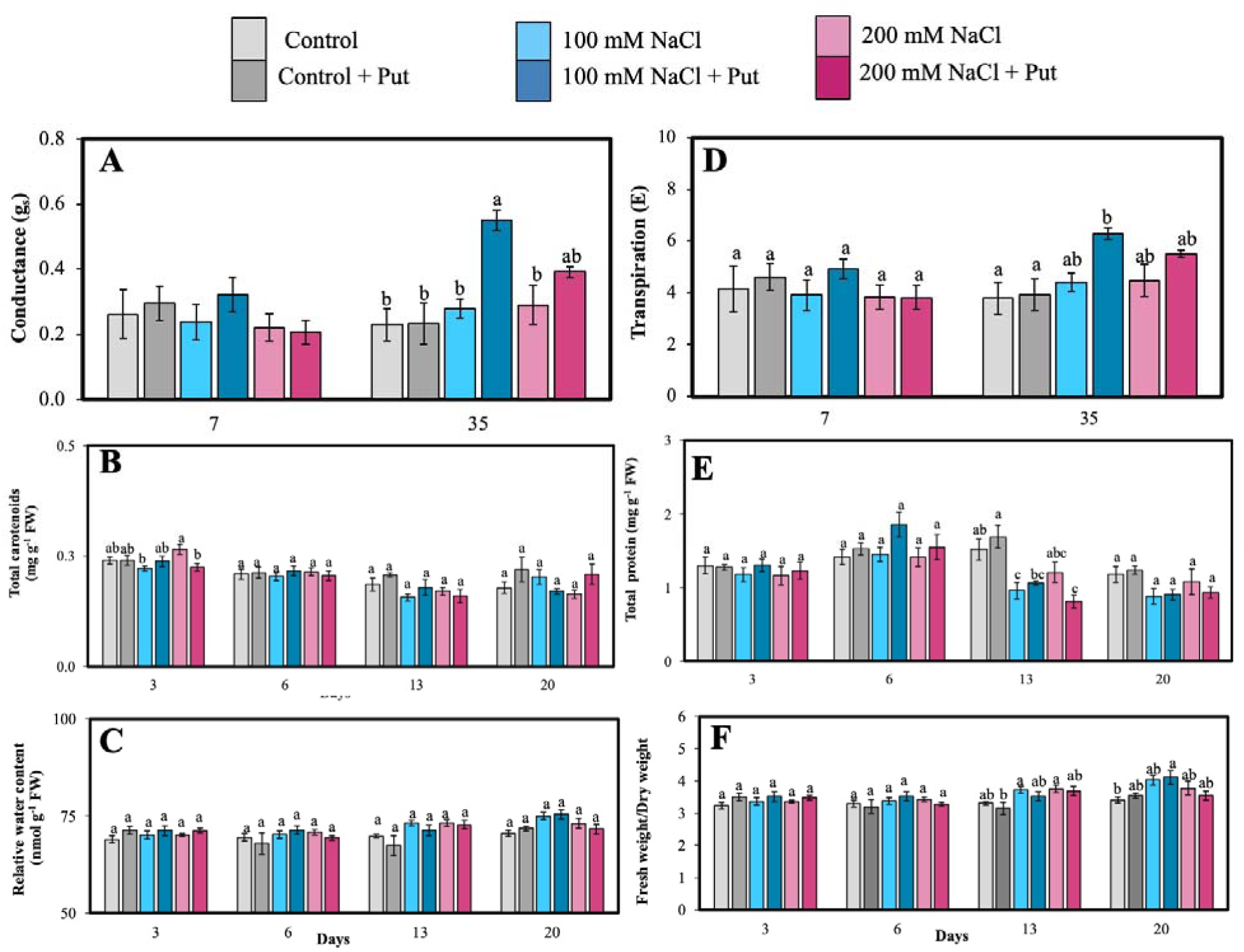
Effect of two different concentrations of NaCl (± Putrescine spray) on physiological accumulation in hybrid poplar NM6 leaves over time. (**A**) Conductance, (**B**) Carotenoid, (**C**) Relative water content, (**D**) Transpiration, (**E**) Protein, and **(F)** Fresh weight/dry weight measured at several days after salt treatment. Different letters indicate statistically significant differences (p < 0.05) among treatments. Data represent mean ± SE (n = 3, 5).

Total carotenoids (Fig. 2B) peaked on day 3 under 200 mM NaCl + Put, reaching 0.39 ± 0.01 mg g ¹ FW, which was significantly higher than 200 mM NaCl alone (0.31 ± 0.01 mg g ¹ FW, *p* < 0.05). Total soluble protein content (Fig. 2E) dropped under salt stress, with the 100 mM NaCl group reaching a low of 2.0 ± 0.08 mg g ¹ FW on day 13, significantly less than the control (2.6 ± 0.07 mg g ¹ FW, *p* < 0.05). Put treatment partially restored protein levels to 2.3 ± 0.09 mg g ¹ FW, indicating a moderate recovery.

Relative water content (RWC) (Fig. 2C) remained stable across treatments and time points, ranging between 71–75%, with no significant differences detected.

The ratio of fresh weight to dry weight (Fig. 2F) was significantly higher in 100 mM NaCl-treated plants on day 13 (3.6 ± 0.15) compared to Put-sprayed control plants (2.9 ± 0.12, *p* < 0.05), possibly indicating osmotic adjustment.

### 3.3 Soluble sugar content

Soluble sugar (fructose, glucose + galactose, and sucrose) was quantified in the 6^th^ fully matured leaf at multiple time points following NaCl and Put treatment. Fructose content significantly increased under Put-sprayed 100 mM NaCl *vs.* unsprayed at 6^th^ day (Fig. 3A). Fructose was significantly higher in 200 mM NaCl treated plants compared to control and 100 mM NaCl treated plants on the 6^th^ day. Glucose and galactose were non-separable in our HPLC system and hence they are represented together. The early and pronounced increase in fructose under severe salt stress may indicate an osmoprotective role. At the same time, Put application appeared to further enhance fructose accumulation even under moderate stress, possibly supporting better stress acclimation. Glucose + galactose levels were significantly lower in 200 mM NaCl treated plants compared to Put sprayed control plants on the 6^th^ and 13^th^ day (Fig. 3B). This reduction suggests hexose depletion under high salinity, likely due to increased utilization for respiration or metabolic adjustments. Put treatment maintained relatively higher levels, indicating a protective effect on hexose stability under salt stress. Sucrose levels were significantly lower in Put-sprayed plants *vs.* unsprayed for 100 mM NaCl on the 6^th^ day (Fig. 3C). Sucrose was significantly higher for Put-sprayed 200 mM NaCl plants on the 13^th^ day. Sucrose was also significantly higher in Put-sprayed plants than those unsprayed for control and 100 mM NaCl on the 13^th^ day. Our results also showed that sucrose contents were also significantly higher in 100 mM NaCl treated plants compared to the control plants on 20^th^ day. These patterns suggest dynamic sucrose reallocation, where early decreases may reflect mobilization for stress response, while later accumulation, especially in Put-treated plants, may signal improved recovery and carbon economy under sustained salt exposure.

**Figure 3:**
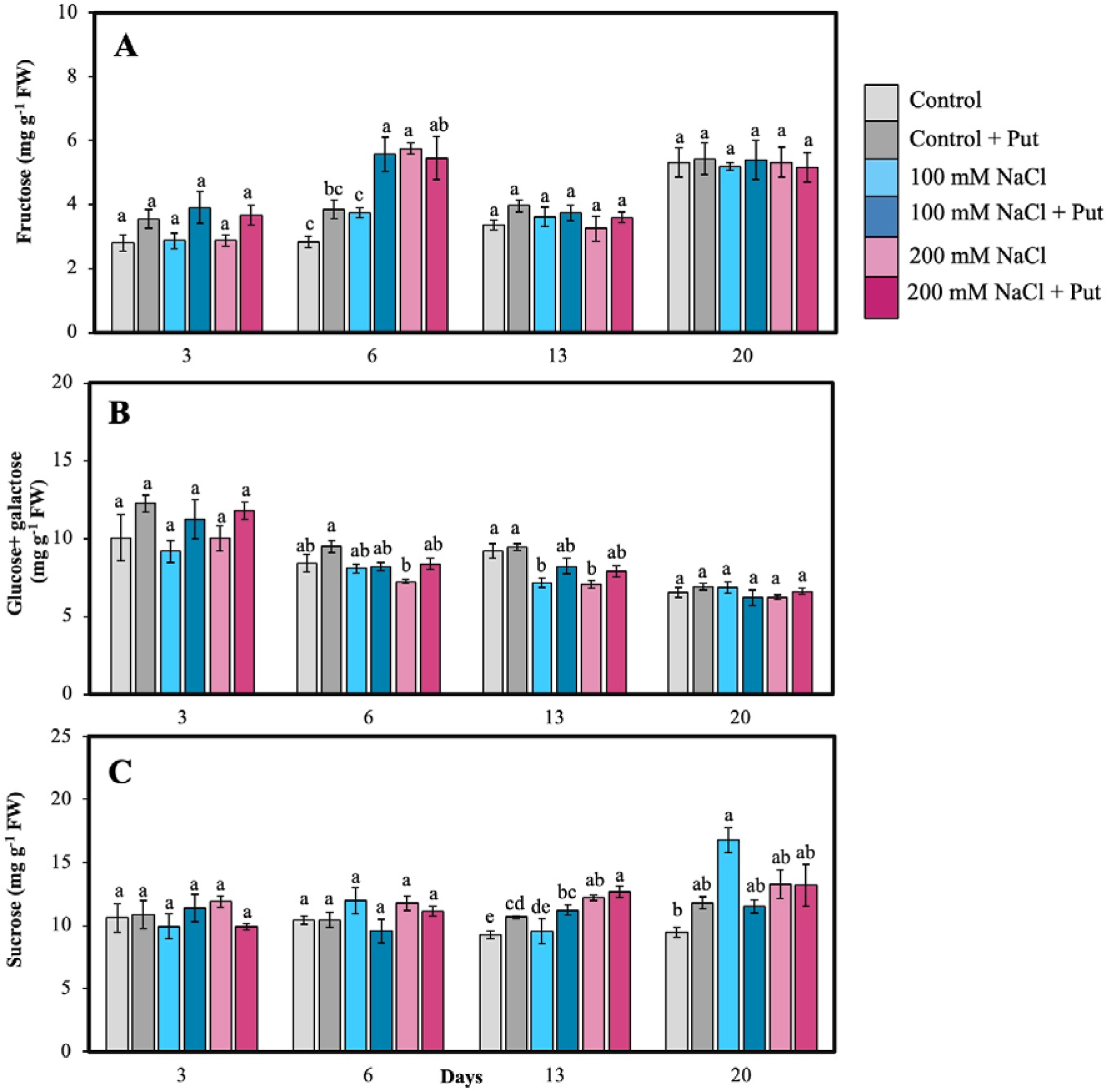
Effect of two different concentrations of NaCl (± Putrescine spray) on soluble sugar accumulation in hybrid poplar NM6 leaves over time. (**A**) Fructose, (**B**) Glucose + Galactose, and (**C**)Sucrose measured at 3, 6, 13, and 20 days after salt treatment. Different letters indicate statistically significant differences (p < 0.05) among treatments. Data represent mean ± SE (n = 5).

The overall temporal shift in sucrose levels may reflect stress-induced carbon partitioning, wherein sucrose is either broken down or synthesized depending on the phase and intensity of salt stress. Early reductions (e.g., day 6 under Put) may indicate rapid mobilization of sucrose for energy production or conversion into hexoses to support osmotic balance. In contrast, late-stage increases (e.g., day 13 in Put-treated plants) may result from enhanced photosynthetic carbon assimilation or reallocation of assimilates toward sucrose storage once stress responses are stabilized. This interpretation aligns with previous observations that Put treatment improved gas exchange (Section 3.2) and mitigated morphological damage (Section 3.1), possibly enabling better carbon economy under salt stress. Collectively, these results support the idea that Put modulates sugar allocation patterns under salinity in a dynamic, sugar-specific, and time-dependent manner.

### 3.4 Amino acids and Polyamines content in leaves and roots

Amino acids and PAs were measured in the 6^th^ fully matured leaf on several days after NaCl treatment. Among all the AAs, Arg + Thr + Gly and Gln were the most dominant. On day 3, 200 mM NaCl treated plants had significantly higher levels of Gln, Ser, and GABA compared to all other treatments (Fig. 4). Put-sprayed plants under 200 mM NaCl treatment, had significantly higher Gln, Ser and Orn on day 6 (Fig. 4A, B, 5A). Additionally, Put-sprayed plants under 100 mM NaCl had significantly higher Glu and Asp compared to control plants on day 6 (Fig 5B, C). Serine was also significantly higher on Put-sprayed, salt treated plants on day 13. Ala, Pro, Arg + Thr + Gly, Phe + Cys, and Ile was significantly higher in unsprayed *vs.* sprayed 200 mM NaCl treated plants on day 3 (Fig. 6A, B, Supplementary Fig. 1, 2). On day 6, Ala was also significantly higher in 100 mM NaCl plants compared to control. Our results also showed that Pro was significantly lower on day 6 in Put-sprayed, 200 mM NaCl-treated plants (Fig. 6B). His was significantly higher in Put-sprayed, 200 mM NaCl-treated plants compared to other treatments on day 6 (Fig. 6B). Foliar spray of Put significantly altered endogenous Put and Spm content under 200 mM NaCl on day 3 (Fig. 7A). However, PA levels varied under salt treatment on other days (Fig. 7B, C). These findings suggest that Put application influences AA and PA accumulation under salt stress in a time-and dose dependent manner.

**Figure 4:**
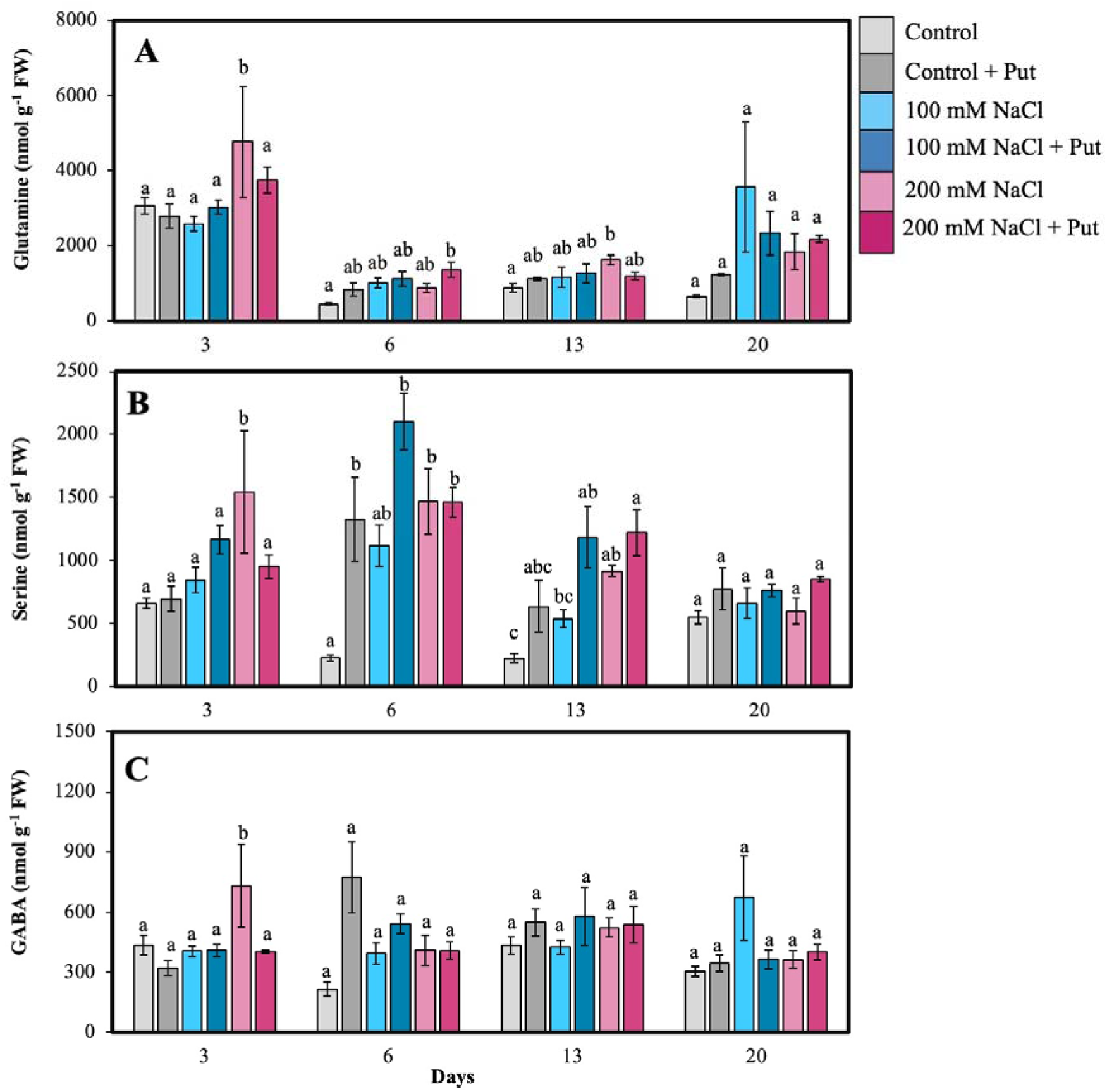
Effect of two different concentrations of NaCl (± Putrescine spray) on amino acids accumulation in hybrid poplar NM6 leaves over time. (**A**) Glutamine, (**B**) Serine, and (**C**) GABA measured at 3, 6, 13, and 20 days after salt treatment. Different letters indicate statistically significant differences (p < 0.05) among treatments. Data represent mean ± SE (n = 5).

**Figure 5:**
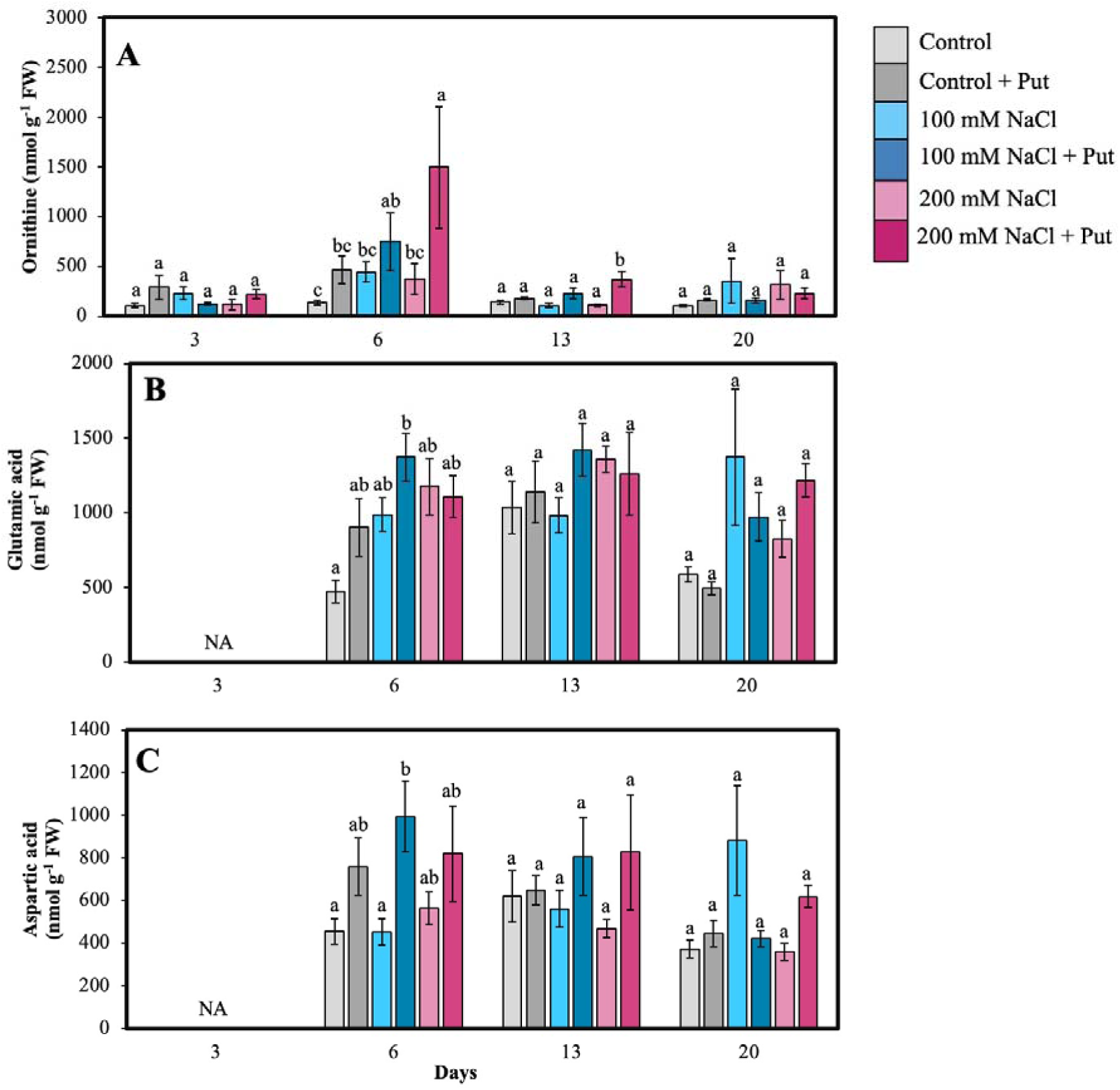
Effect of two different concentrations of NaCl (± Putrescine spray) on amino acids and polyamines accumulation in hybrid poplar NM6 leaves over time. (**A**) Ornithine levels, (**B**) Log Glutamic acid levels, and (**C**) Aspartic acid levels measured at 3, 6, 13, and 20 days after salt treatment. Different letters indicate statistically significant differences (p < 0.05) among treatments. Data represent mean ± SE (n = 5).

**Figure 6:**
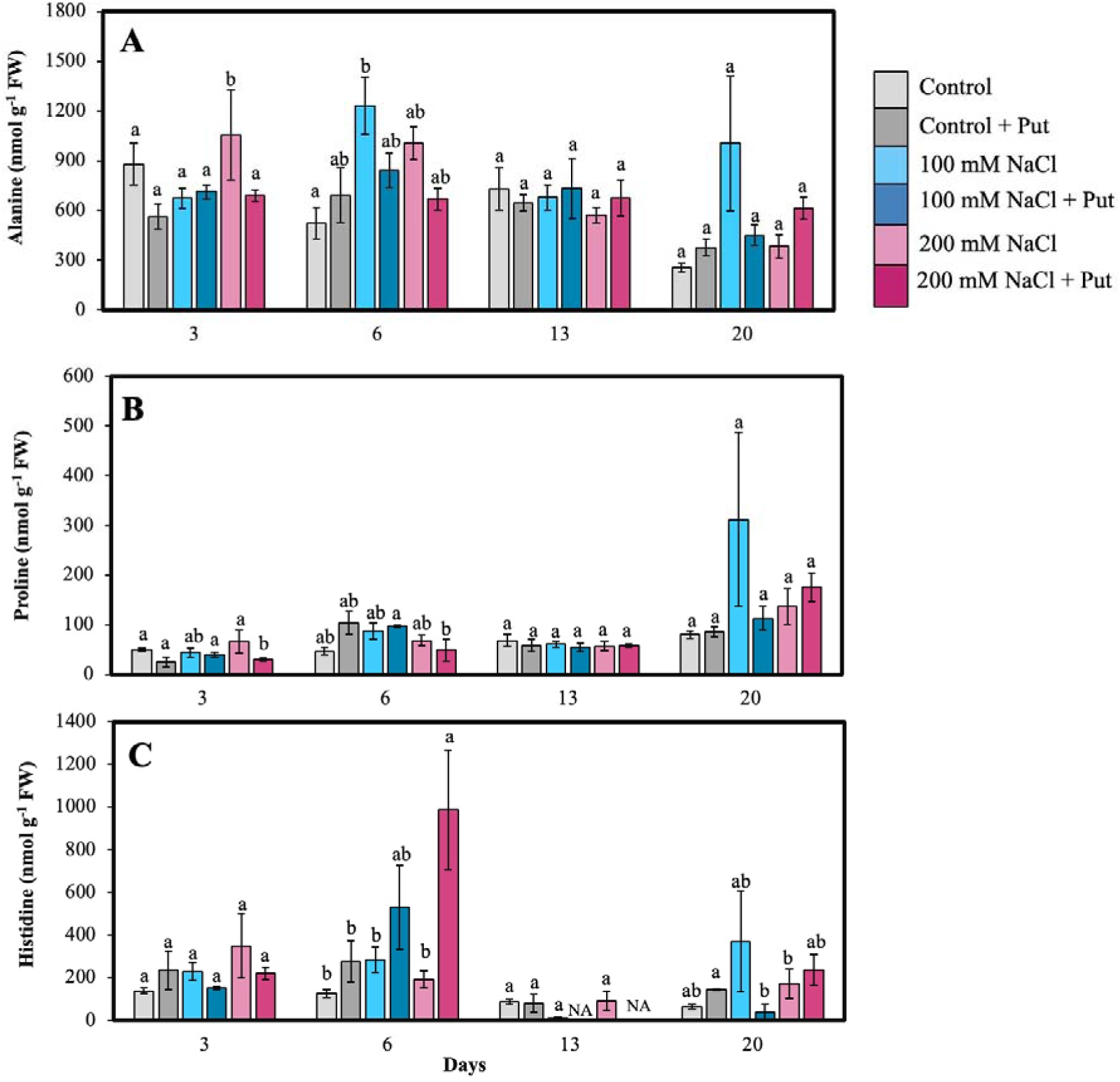
Effect of two different concentrations of NaCl (± Putrescine spray) on amino acids and polyamines accumulation in hybrid poplar NM6 leaves over time. (**A**) Alanine levels, (**B**) Proline levels, and (**C**) Histidine levels measured at 3, 6, 13, and 20 days after salt treatment. Different letters indicate statistically significant differences (p < 0.05) among treatments. Data represent mean ± SE (n = 5).

**Figure 7:**
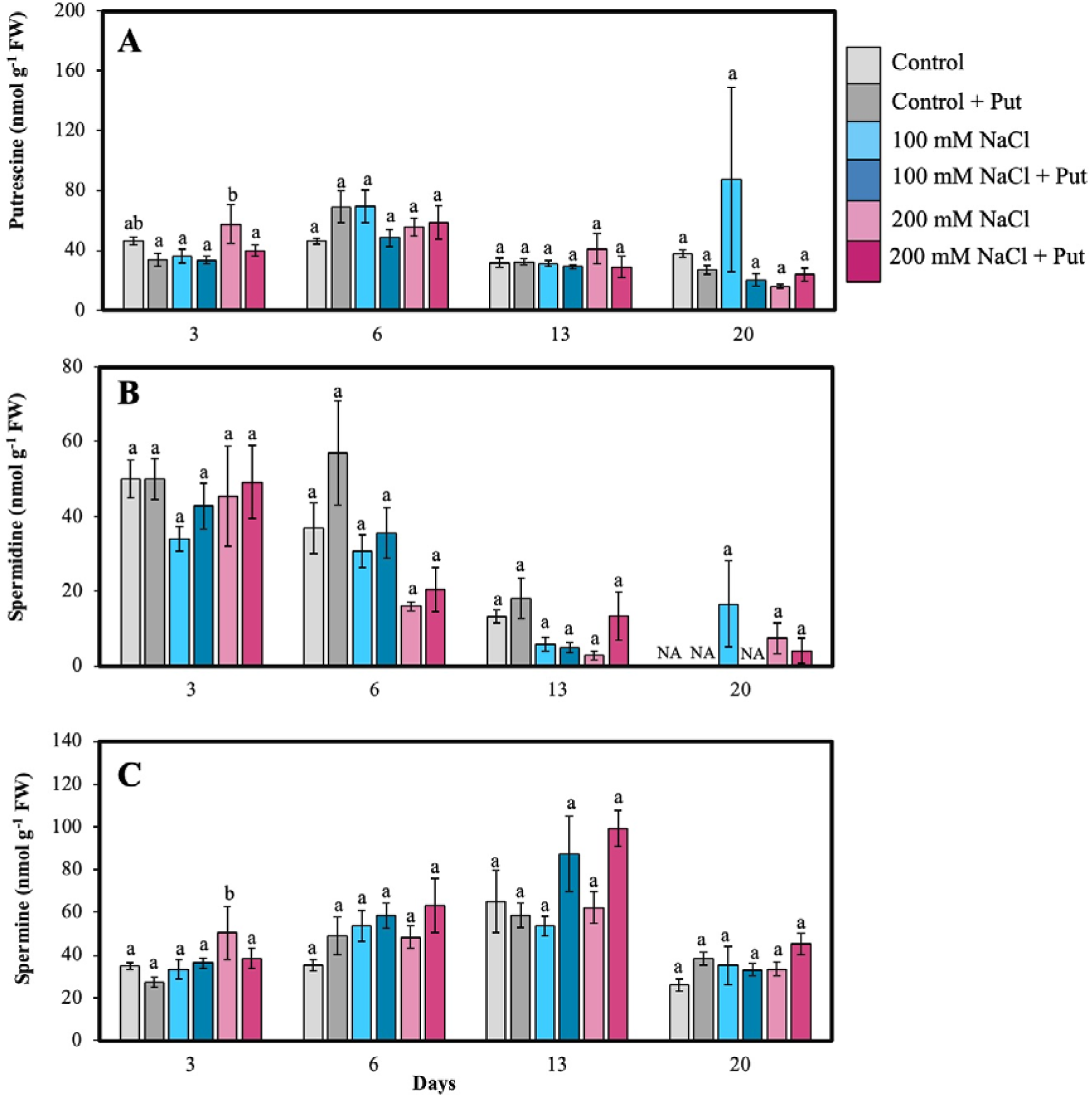
Effect of two different concentrations of NaCl (± Putrescine spray) on amino acids and polyamines accumulation in hybrid poplar NM6 leaves over time. (**A**) Putrescine levels, (**B**) Spermidine levels, and (**C**) Spermine levels measured at 3, 6, 13, and 20 days after salt treatment. Different letters indicate statistically significant differences (p < 0.05) among treatments. Data represent mean ± SE (n = 5).

AAs and PAs content of the roots were analyzed (with HPLC) only at 21 days after NaCl treatment at the termination of the study. Salt significantly increased Arg + Thr + Gly, Gln, Orn, Phe + Cys, and Ser accumulation (in both concentrations of NaCl) compared to the control (Fig. 8A, B). Lysine was significantly lower in 200 mM NaCl compared to control plants (Supplementary Fig 4A). Among PAs, Put was significantly increased under salt stress, while Spd was significantly reduced in salt exposured roots (Fig. 8C). However, Spm was undetected in salt treated roots. Our results also showed that total PA content in roots were lower in 100 mM NaCl-treated plants compared to both the control and 200 mM NaCl-treated plants. These results strongly suggest that salt stress influences AAs and PAs accumulation.

**Figure 8:**
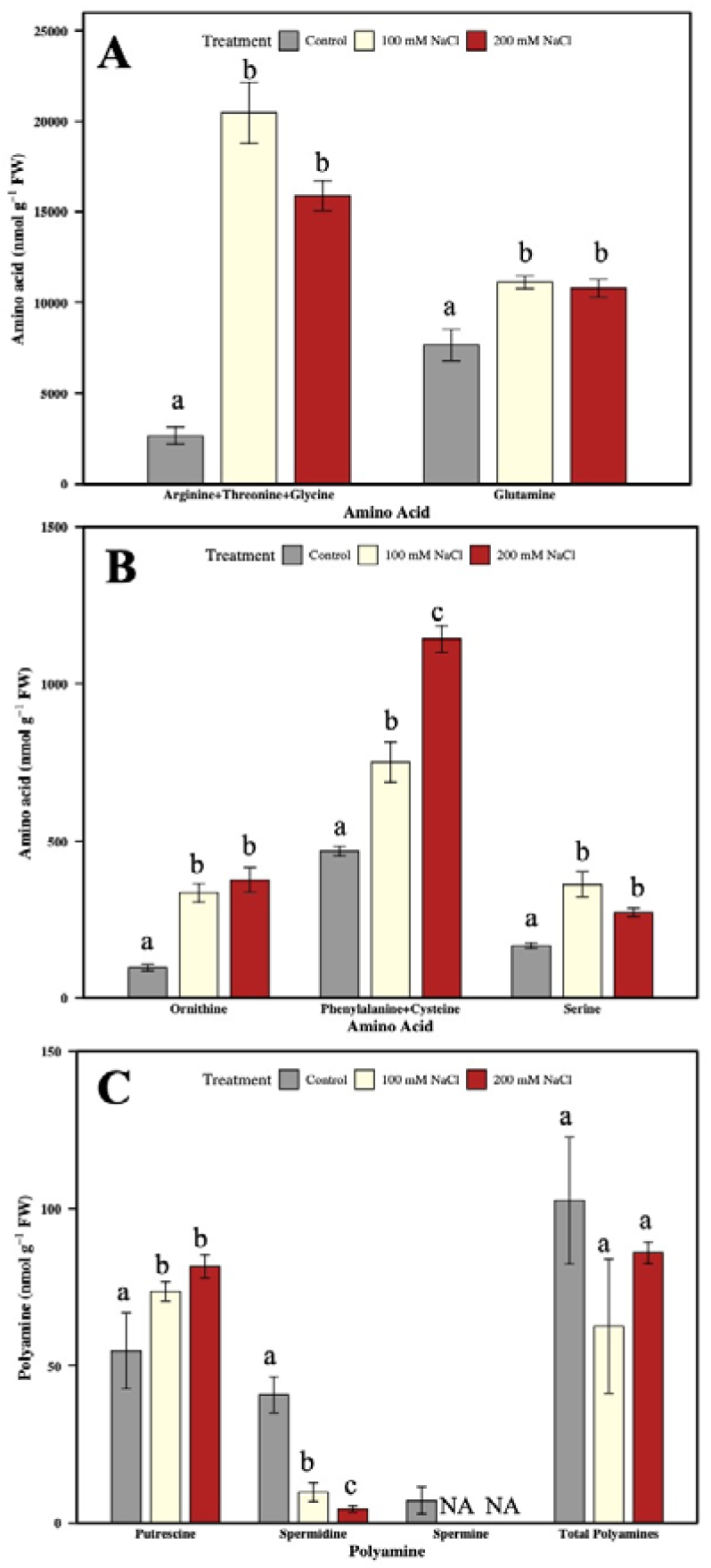
Effect of two different concentrations of NaCl on amino acids and polyamines accumulation in hybrid poplar NM6 roots. (**A, B**) Amino acids levels, (**C**) Polyamines levels measured at 21 days after salt treatment. Different letters indicate statistically significant differences (p < 0.05) among treatments. Data represent mean ± SE (n = 4).

### 3.5 Combined analysis of polyamine metabolic cycle in roots

To assess the metabolic coordination among AAs and PAs in roots under salt stress, we conducted a Pearson correlation analysis based on their relative contents across control, 100 mM NaCl, and 200 mM NaCl treatments (Fig. 9A, B). The heatmap revealed 2 distinct clusters: Cluster 1 included Gln, Pro, Ser, GABA, Put which showed strong positive correlations, several of which were statistically significant (e.g., Gln with Ser, GABA, Arg + Thr + Gly, p< 0.05).

**Figure 9.**
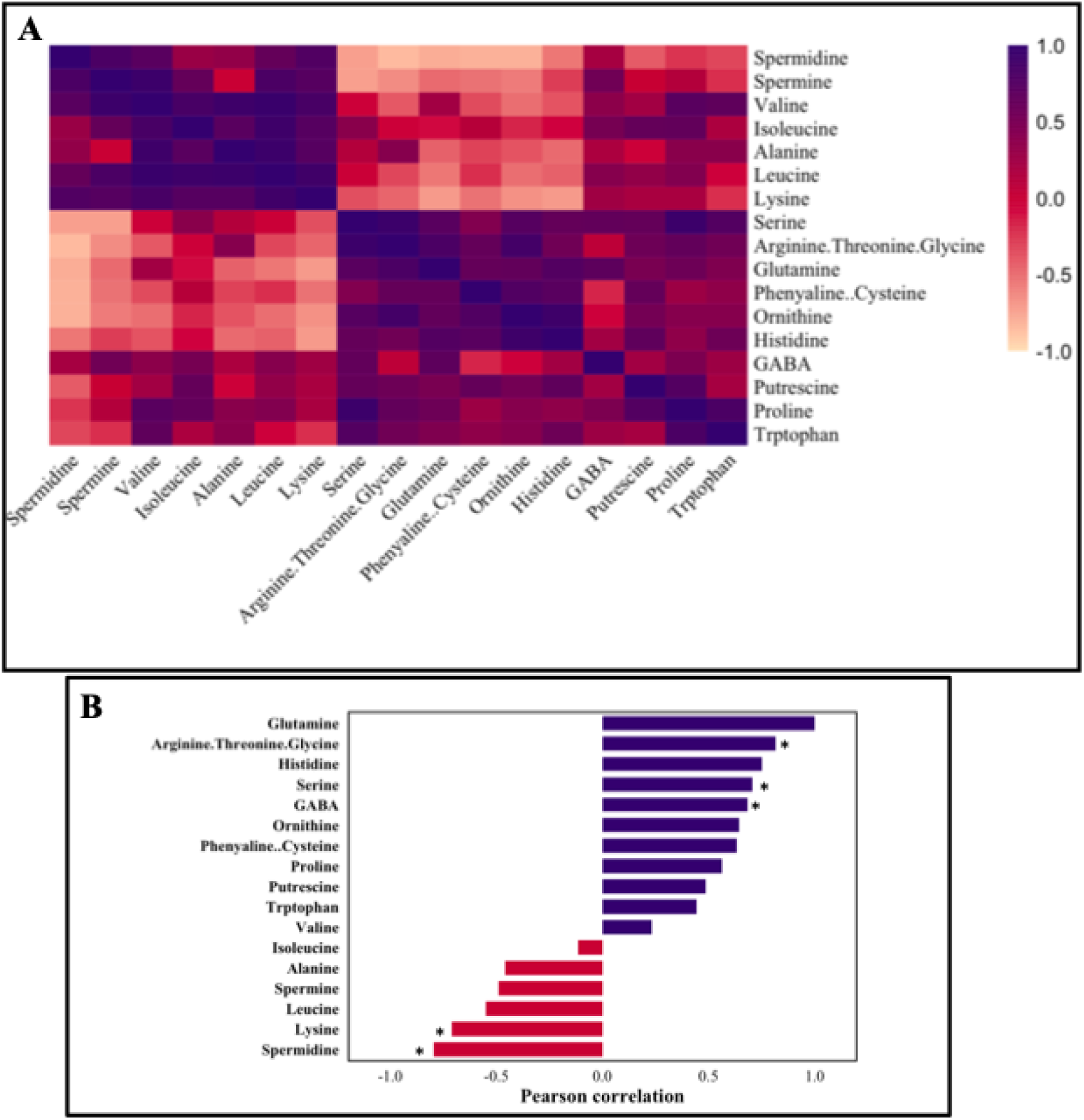
(A) Correlation coefficients of amino acids and polyamines analyzed in the poplar roots treated with two different concentrations of salt. (B) Pattern correlation analysis with Glutamine. The graph reports the significant features detected and ordered according to their correlation coefficient when correlated with Glutamine. The purple color represents a positive correlation, and the red color represents a negative correlation. Correlation distance-Pearson r. Correlation significant at *p < 0.05.

These metabolites are generally associated with stress adaptation and osmoprotection. Cluster 2 was composed of Spd, Spm, Ala, Lys, Leu, Ile, and Val. These showed negative correlations with Cluster 1 metabolites, most notably, Spd and Lys were significantly negatively correlated with Gln. This cluster represents more growth-associated or proteinogenic AAs and PAs.

The distinct separation of these clusters suggests that salt stress induces a shift in root N metabolism, favoring stress-responsive compounds (Cluster 1), while down regulating growth-linked metabolites (Cluster 2).

## 4 Discussion

Soil salinity is a growing global threat to agriculture and forestry, significantly reducing crop yields and limiting the availability of arable land. Salt stress impairs plant function by reducing water uptake, inducing ion toxicity, and triggering oxidative damage (Hasanuzzaman et al., 2013, Dinneny, 2015, Safdar et al., 2019, Sun et al., 2022, Rady et al., 2023). These effects ultimately reduce growth and productivity (Sun et al., 2022, Liu et al., 2023, Rodríguez Coca et al., 2023, Zhang et al., 2023). Typical symptoms include leaf chlorosis and necrosis, stomatal closure, and reduced photosynthetic capacity due to impaired water and nutrient uptake (Hasanuzzaman et al., 2013, de Oliveira et al., 2013, Sun et al., 2022, Calhoun et al., 2023).

Plants respond to salt stress by reprogramming physiological, molecular, and biochemical pathways (Gupta and Huang, 2014, Yang and Guo, 2018, Raza et al., 2022, Wang et al., 2024a, Liu et al., 2025a). Salt responses vary by stress intensity and species (Ding et al., 2010, Nguyen et al., 2018, Zhao et al., 2021, Sun et al., 2022). Ding et al. (2010) reported that salt-tolerant *P. euphratica* Oliv showed greater antioxidant enzyme activity than salt-sensitive *P. popularis 35-44*. Zalesny Jr et al. (2019) found that salt-tolerant *P. nigra X maximowiczii* (NM2 and NM6 hybrid clones) exhibited enhanced mineral uptake under high-salinity conditions. Seed priming or foliar applications of hormones and nitrogenous compounds has been shown to mitigate salinity damage during plant development (Ma et al., 2012, Srivastava et al., 2021, Zafar et al., 2022, Eisa et al., 2023, Zhang et al., 2024).

Polyamines such Put and Spd have been widely studied for their roles in mitigating abiotic stress in crop species (Nahar et al., 2016, Xiong et al., 2018, Alcázar et al., 2020, Islam et al., 2021, Ma et al., 2022, Blázquez, 2024, Zhang et al., 2024). Foliar application of Put has been reported to enhance salt tolerance in several herbaceous species (Supplemental Table 4). However, the role of PAs in salt tolerance remains largely unexplored in woody perennials, particularly in poplar.

In this study, we examined the effects of exogenous Put application on hybrid poplar under salt stress. Our aim was to evaluate its potential as a non-transgenic, physiological approach to enhance stress resilience.

### 4.1 Effect of putrescine on physiological parameters

Salt stress induced visible symptoms in NM6 poplar by day 13, consistent with previous reports. Surprisingly, 200 mM NaCl-treated plants showed milder symptoms, which may reflect early acclimation or altered stress signaling. Put-treated plants (without salt) developed minor tip necrosis but retained darker green foliage. Under combined stress (Put + NaCl), leaf damage shifted to central lamina, suggesting dose-or interaction-dependent effects. Growth inhibition and wilting under salt aligned with responses seen in other species, such as cotton and poplar (Cao et al., 2020, Calhoun et al., 2023, Hu et al., 2023, Zhao et al., 2024a). Put application at 100 mM NaCl enhanced growth, supporting its role as a protective agent under moderate stress.

Previous work shows that abiotic stress can induce long-term physiological adaptation, often described as “stress memory” (Wang et al., 2014, Georgii et al., 2019, Tikhomirova et al., 2022, Liu et al., 2022). Wang et al. (2025) reported that in *Populus alba × glandulosa*, photochemical activity improved in later stages of salt exposure. Consistently, our study showed that Pn in NM6 poplar improved by day 35 under salt stress, relative to earlier time points. Put-treated plants under salt stress also maintained higher gs and E by day 35, similar to trends reported in *C. sinensis* and *C. sativus* (Xiong et al., 2018, Shen et al., 2019). These findings suggest that Put may enhance physiological resilience over time by modulating long-term stress responses and photosynthetic recovery. Total soluble protein declined under salt stress, likely due to increased degradation or inhibited synthesis, potentially contributing to elevated free amino acid levels. Put application under 100 mM NaCl restored protein content, but this effect was absent at 200 mM. No changes were observed in non-stressed plants, indicating the impact of Put is stress-dependent. RWC slightly increased by day 13 under salt treatment, but was unaffected by Put, suggesting that protein fluctuations were not water-driven. FW/DW ratios also rose under salt, possibly reflecting osmotic adjustment.

### 4.2 Effect of putrescine on soluble sugars

Liang et al. (2018), Liu et al. (2022) and Zhang et al. (2024) reported that accumulation of soluble sugars is a common adaptive response to abiotic stress in plants, as reported in *Arabidopsis thaliana*, *Populus euphratica*, *Morus multicaulis* and *Oryza sativa*. Such osmotic adjustment, where sugars functions as key metabolite is well-documented in species like Tartary buckwheat. This trend was reflected in our observations, where early salt exposure corresponded with increased fructose and sucrose levels. Conversely, reduced glucose + galactose levels suggest selective modulation of sugar pathways, potentially favoring disaccharide synthesis or altered carbon allocation. These shifts may be part of a stress-induced self-regulatory mechanism, wherein sugars contribute to osmotic balance, protect membrane stability, and support energy supply for metabolic adaptation. It has been previously established that such osmolyte accumulation also aids in maintaining high intracellular K and favorable Na /K ratios, while preventing dehydration and oxidative damage (Liu et al., 2022, Azeem et al., 2023). Put application further enhanced sugar accumulation under salt stress, indicating its role in modulating C metabolism. This aligns with earlier findings in *Cucumis sativus* by Yuan et al. (2015), where PAs influenced sugar pathways, likely via PA-C metabolism crosstalk. Together, these results suggests that Put enhances sugar-mediated salt tolerance in poplar by promoting osmotic adjustment and stabilizing metabolic functions under stress.

### 4.3 Effect of putrescine on amino acids and polyamines

PAs are small nitrogen-rich compounds known to mitigate salt-induced damage via multiple biochemical and physiological mechanisms (Alcázar et al., 2010, Minocha et al., 2014, Saha et al., 2015, Choudhary et al., 2023). Under salt stress, plants often accumulate high levels of PAs, which help reduce lipid peroxidation, stabilize membrane integrity and ionic homeostasis, and regulate stress-responsive genes (Minocha et al., 2014, Alcázar et al., 2020).

We observed time-dependent changes in Put and Spm in leaves under 100 mM NaCl, suggesting an active role in salt adaptation, while Spd declined over time. In roots, Put accumulation also increased under salt, while Spd declined and Spm was undetectable. These responses are consistent with prior studies showing that PA profiles under salt stress are species-specific and dynamically regulated (Zhang et al., 2020b, Huo et al., 2020, Baniasadi et al., 2018, Ghalati et al., 2020, Shu et al., 2015). Marco et al. (2019) reported Arabidopsis Spm synthase mutants were more salt-sensitive, which further supports the importance of Spm in stress adaptation.

Polyamine catabolism is tightly linked to amino acid (AA) metabolism through shared intermediates and signaling molecules (Kundu, 2023, Blázquez, 2024). PAs also function as stress signals that modulate the accumulation of stress-related AAs such as GABA, Ala, and Pro (Gupta et al., 2013, Marco et al., 2019, Singh and Roychoudhury, 2020, Srivastava et al., 2021, Shao et al., 2022, Verslues et al., 2023). Salt exposure triggered early increases in multiple AAs in the leaves, including Glu, Gln, GABA, and Pro, indicating time-dependent reprogramming of nitrogen metabolism. Put spray modulated this response, further altering AA accumulation patterns under salt. In roots, salt salt stress promoted the accumulation of nitrogen-rich and stress-related AAs, including Gln, Pro, and Orn. The accumulation of these AAs may reflect a metabolic shift supporting osmoprotection and nitrogen storage in poplar under salt stress (Srivastava et al., 2021, Verslues et al., 2023, Wilhelmi et al., 2025). The decline in Lys under 200 mM NaCl may reflect its translocation to shoots, as observed in *C. sinensis* (Hijaz et al., 2018). Gln was positively correlated with GABA, Gly, and Ser, suggesting that increased Gln biosynthesis under salt stress may support synthesis of other AAs. Notably, the interaction between AA metabolism and the TCA cycle may contribute to enhanced salt tolerance, as Ali et al. (2019) suggested, providing a metabolic basis for the coordinated changes observed in our study.

These findings highlight the potential of foliar-applied putrescine as a practical, low-cost strategy to enhance salt tolerance in hybrid poplar, with implications for sustainable forestry and biomass production in saline-prone regions.

## 5 Conclusions

This study demonstrates that *Populus nigra × maximowiczii* (clone NM6) exhibits adaptive physiological and metabolic responses under salt stress, particularly when treated with exogenous Put. At 100 mM NaCl, representing moderate salinity; Put-treated plants exhibited increased growth, conductivity and transpiration rates and notable shifts in soluble protein, sugars, and AAs. These changes suggest enhanced osmotic regulation and metabolic stability, contributing to higher salt stress tolerance in NM6 poplar.

Given these responses under moderate salinity, NM6 shows potential as a viable species for cultivation on marginally saline land specially when supported by low-cost, non-transgenic treatments such as foliar Put application. Due to the unique role of deep-rooting system, rapid growth and soil-binding capacity further supports their use in land restoration, sustainable silviculture, and bioenergy production.

However, the findings are based on controlled greenhouse conditions and short-term responses; field validation under diverse environmental settings is necessary to confirm long-term applicability. With the advent of newer technologies, future studies could incorporate transcriptomic and metabolomic analyses to unravel the full spectrum of PA-mediated regulatiry pathways. Integration of such molecular insights and with the help of artificial intelligence, integrating physiological treatments with genetic improvements could further accelerate the development of stress-resilient NM6 for future saline land management.

## Author Contribution

SK and SC conceived and designed the experiment. SK, MW, and MG collected the samples. SK, analyzed the data, wrote the manuscript. SM supervised the results and analysis. SM supervised and edited the draft manuscript.

## Generative AI statement

The author(s) declare that some Generative AI was used for checking grammatical statement for the creation of this manuscript.

## Supplementary Figures

**Supplementary Figure 1:**
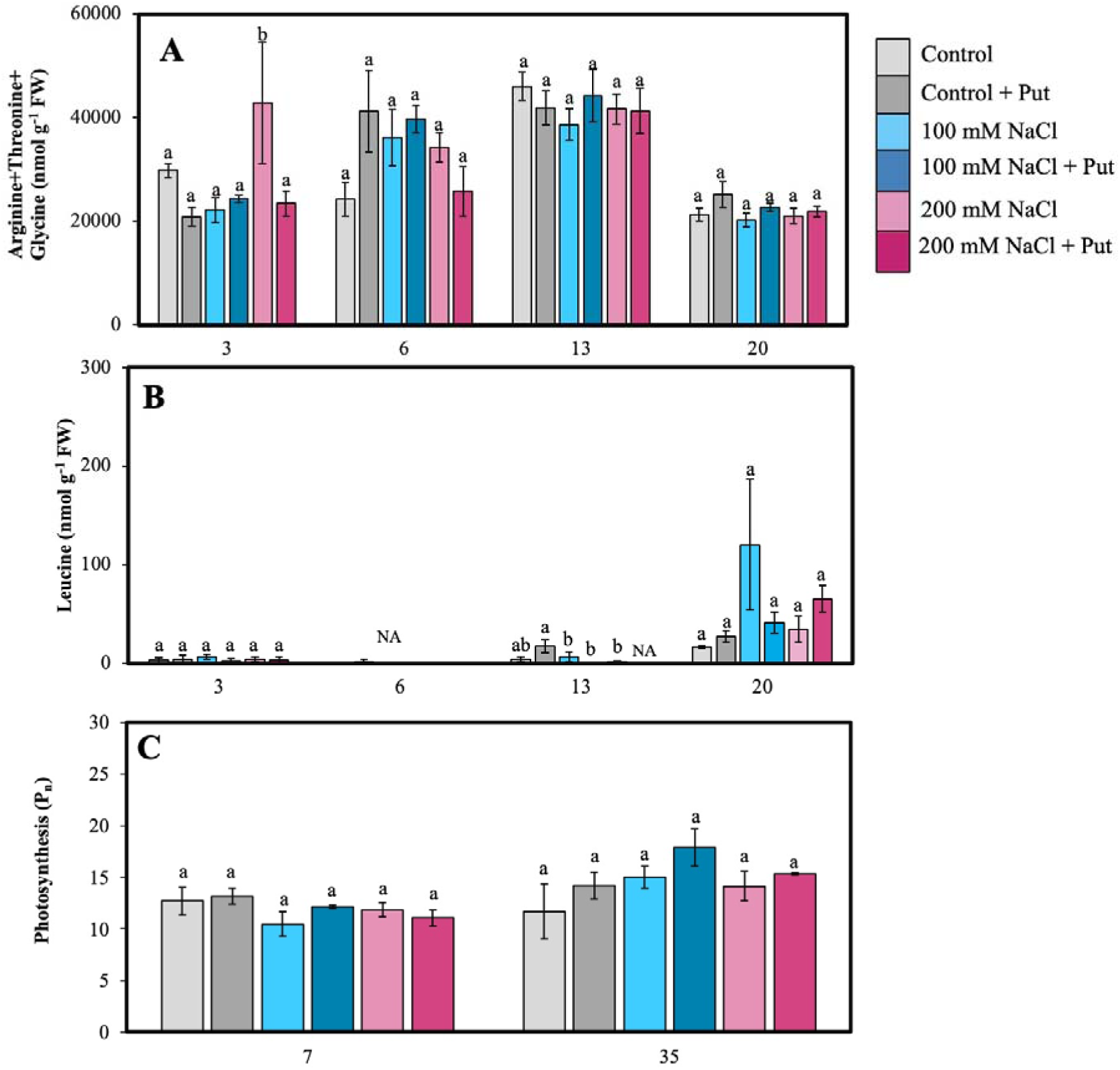
Effect of two different concentrations of NaCl (± Putrescine spray) on amino acids and photosynthesis accumulation in hybrid poplar NM6 leaves over time. (**A**) Arginine+Threonine+Glycine levels, (**B**) Leucine levels, and (**C**) Photosynthesis levels measured at several days after salt treatment. Different letters indicate statistically significant differences (p < 0.05) among treatments. Data represent mean ± SE (n = 5).

**Supplementary Figure 2:**
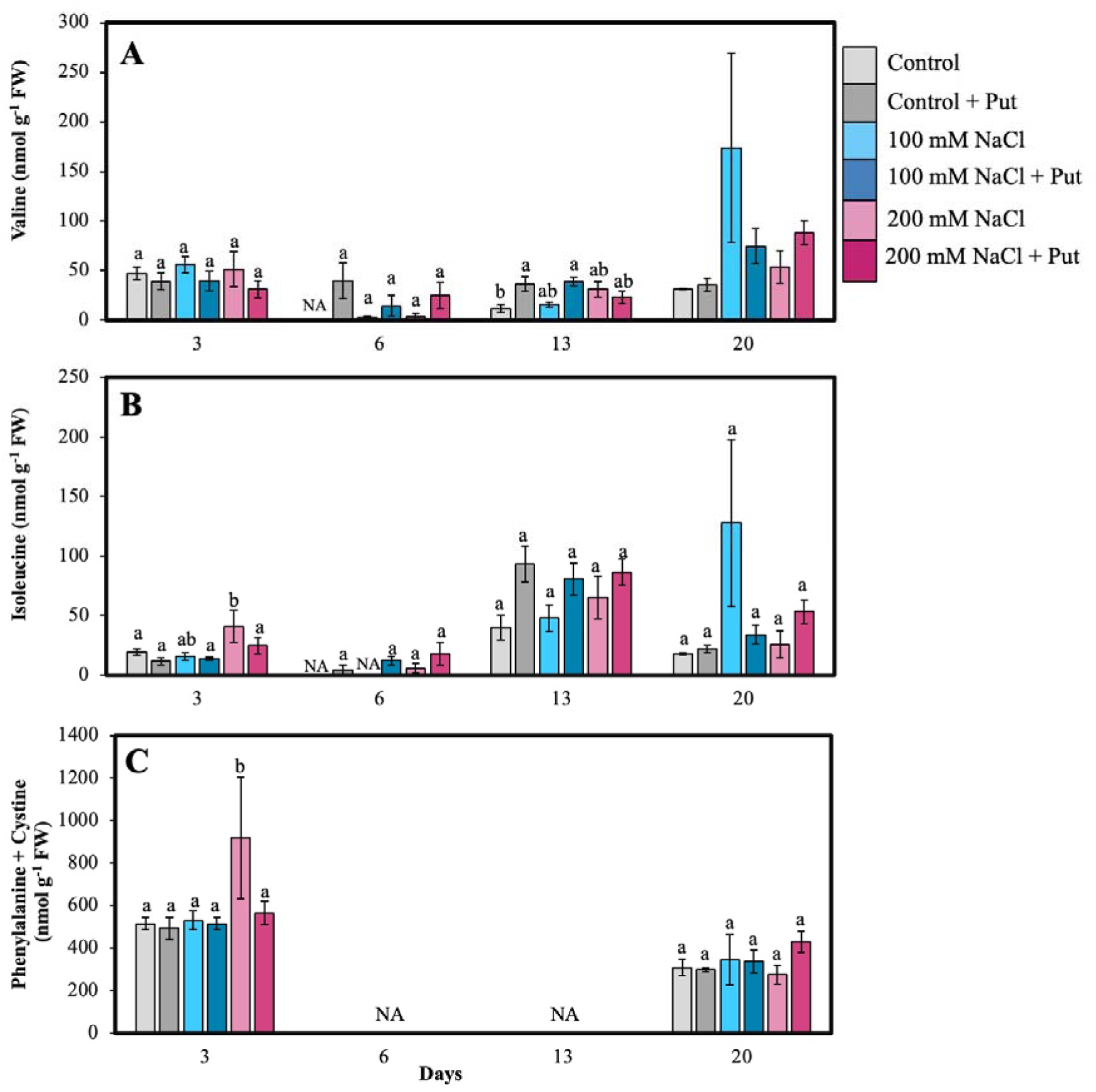
Effect of two different concentrations of NaCl (± Putrescine spray) on amino acids and polyamines accumulation in hybrid poplar NM6 leaves over time. (**A**) Valine levels, and (**B**) Isoleucine levels, and **(C)** Phenylalanine+Cystine levels measured at several days after salt treatment. Different letters indicate statistically significant differences (p < 0.05) among treatments. Data represent mean ± SE (n = 5).

**Supplementary Figure 3:**
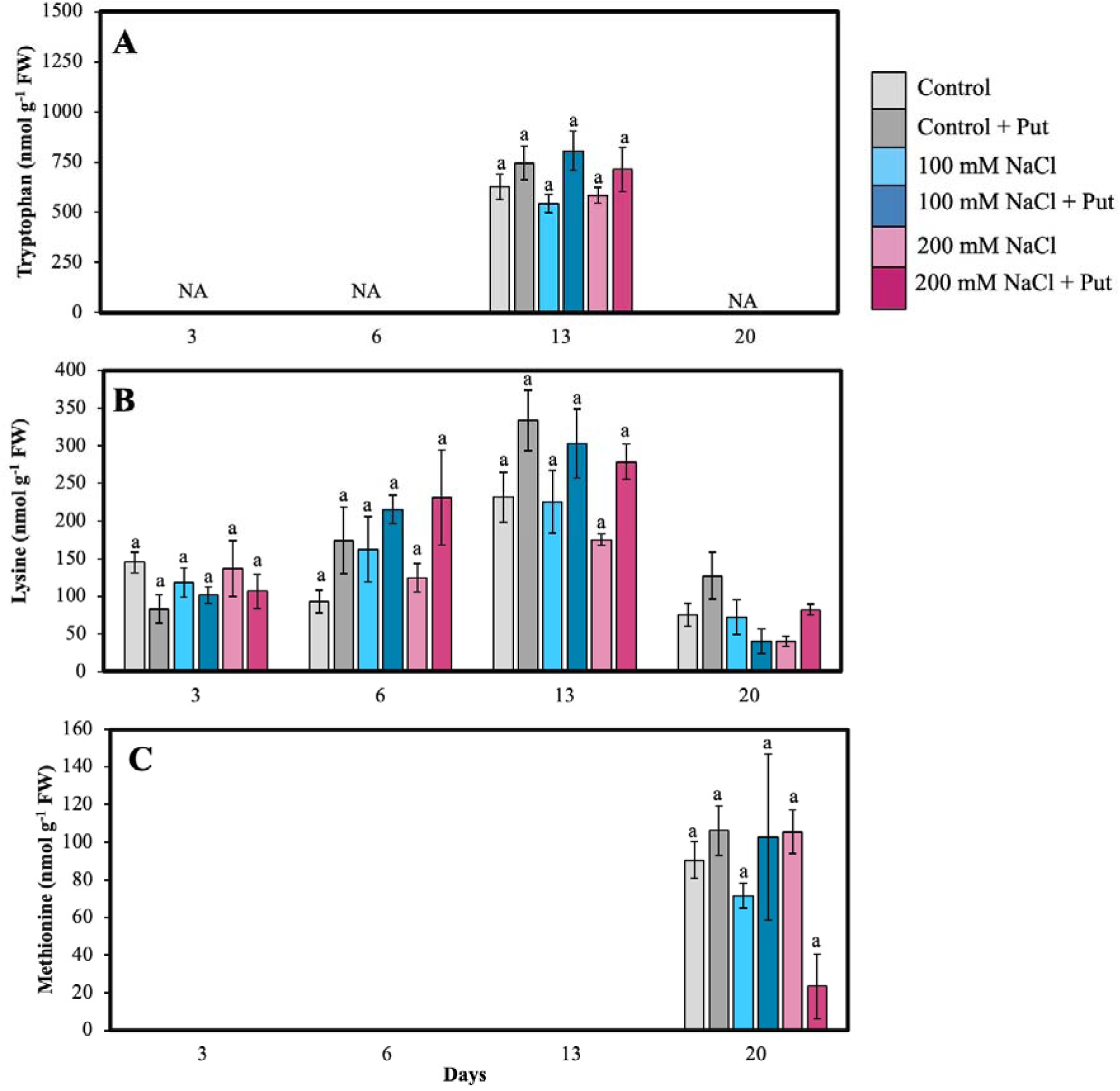
Effect of two different concentrations of NaCl (± Putrescine spray) on amino acids and polyamines accumulation in hybrid poplar NM6 leaves over time. (**A**) Tryptophan levels, (**B**) Lysine levels, and (**C**) Methionine levels measured at several days after salt treatment. Different letters indicate statistically significant differences (p < 0.05) among treatments. Data represent mean ± SE (n = 5).

**Supplementary Figure 4:**
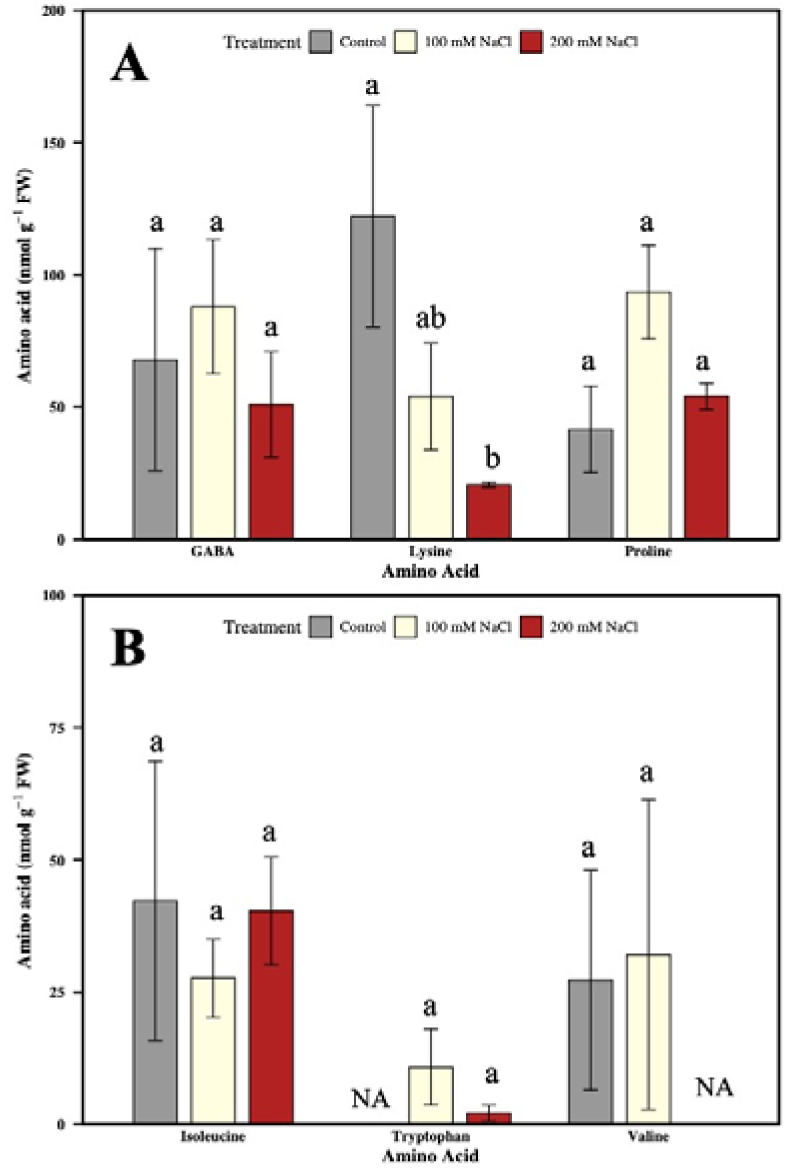
Effect of two different concentrations of NaCl (± Putrescine spray) on amino acids accumulation in hybrid poplar NM6 roots over time. (**A**) GABA, Lysine, Proline levels, and (**B**) Isoleucine, Tryptophan, Valine levels, measured on day 21 after salt treatment. Different letters indicate statistically significant differences (p < 0.05) among treatments. Data represent mean ± SE (n = 4).

**Supplementary Figure 5:**
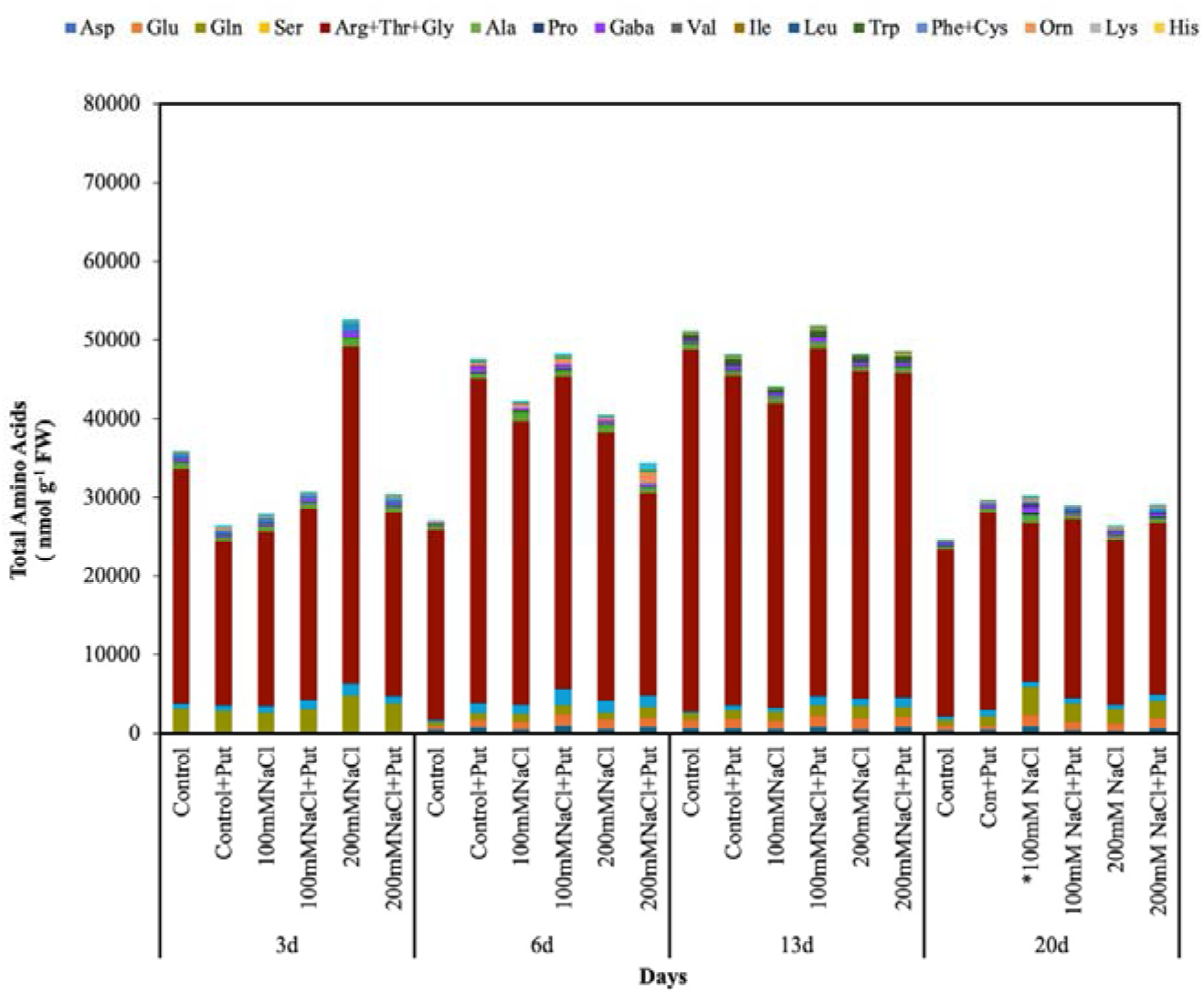
Effect of two concentrations of NaCl (± Putrescine spray) on total amino acid accumulation in hybrid poplar NM6 leaves over time. Stacked bars show the concentrations (nmol g ¹ FW) of individual amino acids measured at 3, 6, 13, and 20 days after salt treatment.

## References

Alcázar, R., Altabella, T., Marco, F., Bortolotti, C., Reymond, M., Koncz, C., Carrasco, P. & Tiburcio, A. F. 2010. Polyamines: molecules with regulatory functions in plant abiotic stress tolerance. Planta, 231, 1237–1249.

Alcázar, R., Bueno, M. & Tiburcio, A. F. 2020. Polyamines: Small amines with large effects on plant abiotic stress tolerance. Cells, 9, 2373.

Alcázar, R., Marco, F., Cuevas, J. C., Patron, M., Ferrando, A., Carrasco, P., Tiburcio, A. F. & Altabella, T. 2006. Involvement of polyamines in plant response to abiotic stress. Biotechnology letters, 28, 1867–1876.

Antoniou, C., Zarza, X., Gohari, G., Panahirad, S., Filippou, P., Tiburcio, A. F. & Fotopoulos, V. 2021. Involvement of polyamine metabolism in the response of Medicago truncatula genotypes to salt stress. Plants, 10, 269.

Atabayeva, S., Minocha, R., Minocha, S., Rakhymgozhina, A., Nabieva, A., Nurmahanova, A., Кenzhebayeva, S., Alybayeva, R. & Asrandina, S. 2020. Response of plants to cadmium stress. International Journal of Biology and Chemistry, 13, 109–117.

Augustyniak, B., Wojtasik, W., Sawuła, A., Burgberger, M. & Kulma, A. 2025. Spermidine treatment limits the development of the fungus in flax shoots by suppressing polyamine metabolism and balanced defence reactions, thus increasing flax resistance to fusariosis. Frontiers in Plant Science, 16, 1561203.

Azeem, M., Pirjan, K., Qasim, M., Mahmood, A., Javed, T., Muhammad, H., Yang, S., Dong, R., Ali, B. & Rahimi, M. 2023. Salinity stress improves antioxidant potential by modulating physio-biochemical responses in Moringa oleifera Lam. Scientific Reports, 13, 1–17.

Azimi, S., Kaur, T. & Gandhi, T. K. 2021. A deep learning approach to measure stress level in plants due to Nitrogen deficiency. Measurement, 173, 108650.

Baniasadi, F., Saffari, V. R. & Moud, A. A. M. 2018. Physiological and growth responses of Calendula officinalis L. plants to the interaction effects of polyamines and salt stress. Scientia Horticulturae, 234, 312–317.

Bargmann, B. O., Laxalt, A. M., Riet, B. T., VAN Schooten, B., Merquiol, E., Testerink, C., Haring, M. A., Bartels, D. & Munnik, T. 2009. Multiple PLDs required for high salinity and water deficit tolerance in plants. Plant and Cell Physiology, 50, 78–89.

Blagden, M., Harrison, J. L., Minocha, R., Sanders-Demott, R., Long, S. & Templer, P. H. 2022. Climate change influences foliar nutrition and metabolism of red maple (Acer rubrum) trees in a northern hardwood forest. Ecosphere, 13, e03859.

Blázquez, M. A. 2024. Polyamines: Their role in plant development and stress. Annual review of plant biology, 75.

Borromeo, I., Domenici, F., DEL Gallo, M. & Forni, C. 2023. Role of polyamines in the response to salt stress of tomato. Plants, 12, 1855.

Buell, C. R., Dardick, C., Parrott, W., Schmitz, R. J., Shih, P. M., Tsai, C.-J. & Urbanowicz, B. 2023. Engineering custom morpho-and chemotypes of Populus for sustainable production of biofuels, bioproducts, and biomaterials. Frontiers in Plant Science, 14, 1288826.

Calhoun, S., Kamel, B., Edmundson, S., Holguin, O., Mach, P., Mckie-Krisberg, Z., Baumgart, L., Blaby, I., Bowen, B. & Chen, C. 2023. A Multi-omic Characterization of the Physiological Responses to Salt Stress in Scenedesmus obliquus UTEX393.

Cao, J.-F., Huang, J.-Q., Liu, X., Huang, C.-C., Zheng, Z.-S., Zhang, X.-F., Shangguan, X.-X., Wang, L.-J., Zhang, Y.-G. & Wendel, J. F. 2020. Genome-wide characterization of the GRF family and their roles in response to salt stress in Gossypium. BMC genomics, 21, 1–16.

Chau, T. N., Timilsena, P. R., Bathala, S. P., Kundu, S., Bargmann, B. O. & Li, S. 2025. Orthologous marker groups reveal broad cell identity conservation across plant single-cell transcriptomes. Nature Communications, 16, 201.

Chen, Y., Tong, S., Jiang, Y., Ai, F., Feng, Y., Zhang, J., Gong, J., Qin, J., Zhang, Y. & Zhu, Y. 2021. Transcriptional landscape of highly lignified poplar stems at single-cell resolution. Genome biology, 22, 1–22.

Choudhary, S., Wani, K. I., Naeem, M., Khan, M. M. A. & Aftab, T. 2023. Cellular responses, osmotic adjustments, and role of osmolytes in providing salt stress resilience in higher plants: polyamines and nitric oxide crosstalk. Journal of Plant Growth Regulation, 42, 539–553.

Conde, D., Triozzi, P. M., Pereira, W. J., Schmidt, H. W., Balmant, K. M., Knaack, S. A., Redondo-López, A., Roy, S., Dervinis, C. & Kirst, M. 2022. Single-nuclei transcriptome analysis of the shoot apex vascular system differentiation in Populus. Development, 149, dev200632.

Cramer, G. R., Urano, K., Delrot, S., Pezzotti, M. & Shinozaki, K. 2011. Effects of abiotic stress on plants: a systems biology perspective. BMC plant biology, 11, 1–14.

De Oliveira, A. B., Alencar, N. L. M. & Gomes-Filho, E. 2013. Comparison between the water and salt stress effects on plant growth and development. Responses of organisms to water stress, 4, 67–94.

Dickmann, D. 2001. Poplar culture in north America, NRC Research Press.

Ding, M., Hou, P., Shen, X., Wang, M., Deng, S., Sun, J., Xiao, F., Wang, R., Zhou, X. & Lu, C. 2010. Salt-induced expression of genes related to Na+/K+ and ROS homeostasis in leaves of salt-resistant and salt-sensitive poplar species. Plant molecular biology, 73, 251–269.

Dinneny, J. R. 2015. Traversing organizational scales in plant salt-stress responses. Current Opinion in Plant Biology, 23, 70–75.

Eisa, E. A., Honfi, P., Tilly-Mándy, A. & Mirmazloum, I. 2023. Exogenous Melatonin Application Induced Morpho-Physiological and Biochemical Regulations Conferring Salt Tolerance in Ranunculus asiaticus L. Horticulturae, 9, 228.

Elsayed, A. I., Mohamed, A. H., Rafudeen, M. S., Omar, A. A., Awad, M. F. & Mansour, E. 2022. Polyamines mitigate the destructive impacts of salinity stress by enhancing photosynthetic capacity, antioxidant defense system and upregulation of calvin cycle-related genes in rapeseed (Brassica napus L.). Saudi Journal of Biological Sciences, 29, 3675–3686.

Georgii, E., Kugler, K., Pfeifer, M., Vanzo, E., Block, K., Domagalska, M. A., Jud, W., Abdelgawad, H., Asard, H. & Reinhardt, R. 2019. The systems architecture of molecular memory in poplar after abiotic stress. The Plant Cell, 31, 346–367.

Ghalati, R. E., Shamili, M. & Homaei, A. 2020. Effect of putrescine on biochemical and physiological characteristics of guava (Psidium guajava L.) seedlings under salt stress. Scientia Horticulturae, 261, 108961.

Ghezehei, S. B., Ewald, A. L., Hazel, D. W., Zalesny Jr, R. S. & Nichols, E. G. 2021. Productivity and profitability of poplars on fertile and marginal sandy soils under different density and fertilization treatments. Forests, 12, 869.

Guerra, F., Gainza, F., Pérez, R. & Zamudio, F. 2011. Phytoremediation of heavy metals using poplars (Populus spp.): a glimpse of the plant responses to copper, cadmium and zinc stress. *Handbook of phytoremediation. Nova Science*, New York, 387-413.

Gupta, B. & Huang, B. 2014. Mechanism of salinity tolerance in plants: physiological, biochemical, and molecular characterization. International journal of genomics, 2014.

Gupta, K., Dey, A. & Gupta, B. 2013. Plant polyamines in abiotic stress responses. Acta physiologiae plantarum, 35, 2015–2036.

Haak, D. C., Fukao, T., Grene, R., Hua, Z., Ivanov, R., Perrella, G. & Li, S. 2017. Multilevel regulation of abiotic stress responses in plants. Frontiers in plant science, 8, 1564.

Hanson, A. D., Beaudoin, G. A., Mccarty, D. R. & Gregory Iii, J. F. 2016. Does abiotic stress cause functional B vitamin deficiency in plants? Plant Physiology, 172, 2082–2097.

Hasanuzzaman, M., Nahar, K. & Fujita, M. 2013. Plant response to salt stress and role of exogenous protectants to mitigate salt-induced damages. Ecophysiology and responses of plants under salt stress, 25-87.

Hawkins, S., Leplé, J.-C., Cornu, D., Jouanin, L. & Pilate, G. 2003. Stability of transgene expression in poplar: a model forest tree species. Annals of forest science, 60, 427–438.

Hijaz, F., Nehela, Y. & Killiny, N. 2018. Application of gamma-aminobutyric acid increased the level of phytohormones in Citrus sinensis. Planta, 248, 909–918.

Hu, J., Zou, S., Huang, J., Huan, X., Jin, X., Zhou, L., Zhao, K., Han, Y. & Wang, S. 2023. PagMYB151 facilitates proline accumulation to enhance salt tolerance of poplar. BMC genomics, 24, 345.

Huo, L., Guo, Z., Wang, P., Zhang, Z., Jia, X., Sun, Y., Sun, X., Gong, X. & Ma, F. 2020. MdATG8i functions positively in apple salt tolerance by maintaining photosynthetic ability and increasing the accumulation of arginine and polyamines. Environmental and Experimental Botany, 172, 103989.

Imran, M., Aaqil khan, M., Shahzad, R., Bilal, S., Khan, M., Yun, B.-W., Khan, A. L. & Lee, I.-J. 2021. Melatonin ameliorates thermotolerance in soybean seedling through balancing redox homeostasis and modulating antioxidant defense, phytohormones and polyamines biosynthesis. Molecules, 26, 5116.

Islam, M. J., Ryu, B. R., Azad, M. O. K., Rahman, M. H., Rana, M. S., Lim, J.-D. & Lim, Y.-S. 2021. Exogenous putrescine enhances salt tolerance and ginsenosides content in Korean ginseng (Panax ginseng Meyer) sprouts. Plants, 10, 1313.

Kundu, S. 2023. Physiological and Biochemical Responses to Salt and Cadmium Stress and It’s Amelioration by Exogenous Application of Polyamines in Hybrid Poplar (Populus n igra x maximowiczii, Clone NM6), University of New Hampshire.

Labrecque, M. & Teodorescu, T. I. 2005. Field performance and biomass production of 12 willow and poplar clones in short-rotation coppice in southern Quebec (Canada). Biomass and Bioenergy, 29, 1–9.

Li, M., Petrie, M. D., Tariq, A. & Zeng, F. 2021. Response of nodulation, nitrogen fixation to salt stress in a desert legume Alhagi sparsifolia. Environmental and Experimental Botany, 183, 104348.

Li, P., Liu, Q., Wei, Y., Xing, C., Xu, Z., Ding, F., Liu, Y., Lu, Q., Hu, N. & Wang, T. 2024. Transcriptional landscape of cotton roots in response to salt stress at single-cell resolution. Plant Communications, 5.

Liang, W., Ma, X., Wan, P. & Liu, L. 2018. Plant salt-tolerance mechanism: A review. Biochemical and biophysical research communications, 495, 286–291.

Lichtenthaler, H. K. 1987. [34] Chlorophylls and carotenoids: pigments of photosynthetic biomembranes. Methods in enzymology. Elsevier.

Lin, T., Lu, Q., Zheng, Z., Li, S., Li, S., Liu, Y., Zhu, T., Chen, L., Yang, C. & Han, S. 2023. Soil cadmium stress affects the phyllosphere microbiome and associated pathogen resistance differently in male and female poplars. Journal of Experimental Botany, erad034.

Liu, H., Todd, J. L. & Luo, H. 2023. Turfgrass Salinity Stress and Tolerance—A Review. Plants, 12, 925.

Liu, L., Cao, X., Zhai, Z., Ma, S., Tian, Y. & Cheng, J. 2022. Direct evidence of drought stress memory in mulberry from a physiological perspective: Antioxidative, osmotic and phytohormonal regulations. Plant Physiology and Biochemistry, 186, 76–87.

Liu, Q., Kang, J., Du, L., Liu, Z., Liang, H., Wang, K., He, H., Zhang, X., Wang, Q. & Hong, Y. Single-cell multiome reveals root hair-specific responses to salt stress. New Phytologist.

Liu, Q., Kang, J., Du, L., Liu, Z., Liang, H., Wang, K., He, H., Zhang, X., Wang, Q. & Hong, Y. 2025a. Single-cell multiome reveals root hair-specific responses to salt stress. New Phytologist.

Liu, T., Qu, J., Fang, Y., Yang, H., Lai, W., Pan, L. & Liu, J. H. 2025b. Polyamines: The valuable bio-stimulants and endogenous signaling molecules for plant development and stress response. Journal of Integrative Plant Biology, 67, 582–595.

Ma, J., Wan, D., Duan, B., Bai, X., Bai, Q., Chen, N. & Ma, T. 2019. Genome sequence and genetic transformation of a widely distributed and cultivated poplar. Plant biotechnology journal, 17, 451–460.

Ma, L., Li, Y., Yu, C., Wang, Y., Li, X., Li, N., Chen, Q. & Bu, N. 2012. Alleviation of exogenous oligochitosan on wheat seedlings growth under salt stress. Protoplasma, 249, 393–399.

Ma, S., Zhou, X., Jahan, M. S., Guo, S., Tian, M., Zhou, R., Liu, H., Feng, B. & Shu, S. 2022. Putrescine regulates stomatal opening of cucumber leaves under salt stress via the H2O2-mediated signaling pathway. Plant Physiology and Biochemistry, 170, 87–97.

Majumdar, R., Shao, L., Minocha, R., Long, S. & Minocha, S. C. 2013. Ornithine: the overlooked molecule in the regulation of polyamine metabolism3. Plant and Cell Physiology, 54, 990–1004.

Malik, A., Yadav, P. & Singh, S. 2022. Role of polyamines in heavy metal stressed plants. Plant Physiology Reports, 1-15.

Marco, F., Busó, E., Lafuente, T. & Carrasco, P. 2019. Spermine confers stress resilience by modulating abscisic acid biosynthesis and stress responses in Arabidopsis plants. Frontiers in Plant Science, 10, 972.

Mcdermot, C. R., Minocha, R., D’AMICO Iii, V., Long, S. & Trammell, T. L. 2020. Red maple (Acer rubrum L.) trees demonstrate acclimation to urban conditions in deciduous forests embedded in cities. PLoS one, 15, e0236313.

Minocha, R. & Long, S. 2004. Simultaneous separation and quantitation of amino acids and polyamines of forest tree tissues and cell cultures within a single high-performance liquid chromatography run using dansyl derivatization. Journal of Chromatography A, 1035, 63–73.

Minocha, R., Long, S., Turlapati, S. A. & Fernandez, I. 2019. Dynamic species-specific metabolic changes in the trees exposed to chronic N+ S additions at the Bear Brook Watershed in Maine, USA. Annals of Forest Science, 76, 1–13.

Minocha, R., Majumdar, R. & Minocha, S. C. 2014. Polyamines and abiotic stress in plants: a complex relationship. Frontiers in plant science, 5, 175.

Minocha, R., Shortle, W. C., Long, S. L. & Minocha, S. C. 1994. A rapid and reliable procedure for extraction of cellular polyamines and inorganic ions from plant tissues. Journal of Plant Growth Regulation, 13, 187–193.

Mohapatra, S., Cherry, S., Minocha, R., Majumdar, R., Thangavel, P., Long, S. & Minocha, S. C. 2010. The response of high and low polyamine-producing cell lines to aluminum and calcium stress. Plant Physiology and Biochemistry, 48, 612–620.

Nahar, K., Hasanuzzaman, M., Rahman, A., Alam, M. M., Mahmud, J.-A., Suzuki, T. & Fujita, M. 2016. Polyamines confer salt tolerance in mung bean (Vigna radiata L.) by reducing sodium uptake, improving nutrient homeostasis, antioxidant defense, and methylglyoxal detoxification systems. Frontiers in Plant Science, 7, 1104.

Neale, D. B. & Kremer, A. 2011. Forest tree genomics: growing resources and applications. Nature Reviews Genetics, 12, 111–122.

Nguyen, H. C., Lin, K. H., Ho, S. L., Chiang, C. M. & Yang, C. M. 2018. Enhancing the abiotic stress tolerance of plants: from chemical treatment to biotechnological approaches. Physiologia plantarum, 164, 452–466.

Pál, M., Rahman, A., Hamow, K. Á., Nagy, K., Janda, T., Dernovics, M. & Szalai, G. 2025. Genotype-specific and light dependence of polyamine uptake and metabolism in wheat plants. Plant Physiology and Biochemistry, 109659.

Pál, M., Szalai, G., Gondor, O. K. & Janda, T. 2021. Unfinished story of polyamines: Role of conjugation, transport and light-related regulation in the polyamine metabolism in plants. Plant Science, 308, 110923.

Pál, M., Tajti, J., Szalai, G., Peeva, V., Végh, B. & Janda, T. 2018. Interaction of polyamines, abscisic acid and proline under osmotic stress in the leaves of wheat plants. Scientific reports, 8, 12839.

Parihar, P., Singh, S., Singh, R., Singh, V. P. & Prasad, S. M. 2015. Effect of salinity stress on plants and its tolerance strategies: a review. Environmental science and pollution research, 22, 4056–4075.

Paul, S., Banerjee, A. & Roychoudhury, A. 2018. Role of polyamines in mediating antioxidant defense and epigenetic regulation in plants exposed to heavy metal toxicity. *Plants Under Metal and Metalloid Stress: Responses*, Tolerance and Remediation, 229-247.

Plomion, C., Bastien, C., Bogeat-Triboulot, M.-B., Bouffier, L., Déjardin, A., Duplessis, S., Fady, B., Heuertz, M., Le Gac, A.-L. & Le Provost, G. 2016. Forest tree genomics: 10 achievements from the past 10 years and future prospects. Annals of Forest Science, 73, 77–103.

Rady, M. M., Mossa, A.-T. H., Youssof, A. M., Osman, A. S., Ahmed, S. M. & Mohamed, I. A. 2023. Exploring the reinforcing effect of nano-potassium on the antioxidant defense system reflecting the increased yield and quality of salt-stressed squash plants. Scientia Horticulturae, 308, 111609.

Raza, A., Tabassum, J., Fakhar, A. Z., Sharif, R., Chen, H., Zhang, C., Ju, L., Fotopoulos, V., Siddique, K. H. & Singh, R. K. 2022. Smart reprograming of plants against salinity stress using modern biotechnological tools. Critical Reviews in Biotechnology, 1-28.

Rodríguez Coca, L. I., García González, M. T., Gil Unday, Z., Jiménez Hernández, J., Rodríguez Jáuregui, M.M. & Fernández Cancio, Y. 2023. Effects of Sodium Salinity on Rice (Oryza sativa L.) Cultivation: A Review. Sustainability, 15, 1804.

Safdar, H., Amin, A., Shafiq, Y., Ali, A., Yasin, R., Shoukat, A., Hussan, M. U. & Sarwar, M. I. 2019. A review: Impact of salinity on plant growth. Nat. Sci, 17, 34–40.

Saha, J., Brauer, E. K., Sengupta, A., Popescu, S. C., Gupta, K. & Gupta, B. 2015. Polyamines as redox homeostasis regulators during salt stress in plants. Frontiers in Environmental Science, 3, 21.

Shao, J., Huang, K., Batool, M., Idrees, F., Afzal, R., Haroon, M., Noushahi, H. A., Wu, W., Hu, Q. & Lu, X. 2022. Versatile roles of polyamines in improving abiotic stress tolerance of plants. Frontiers in Plant Science, 13, 1003155.

Shen, J.-L., Wang, Y., Shu, S., Jahan, M. S., Zhong, M., Wu, J.-Q., Sun, J. & Guo, S.-R. 2019. Exogenous putrescine regulates leaf starch overaccumulation in cucumber under salt stress. Scientia horticulturae, 253, 99–110.

Sheteiwy, M. S., Shao, H., Qi, W., Daly, P., Sharma, A., Shaghaleh, H., Hamoud, Y. A., El-Esawi, M. A., Pan, R. & Wan, Q. 2021. Seed priming and foliar application with jasmonic acid enhance salinity stress tolerance of soybean (Glycine max L.) seedlings. Journal of the Science of Food and Agriculture, 101, 2027–2041.

Shu, S., Yuan, Y., Chen, J., Sun, J., Zhang, W., Tang, Y., Zhong, M. & Guo, S. 2015. The role of putrescine in the regulation of proteins and fatty acids of thylakoid membranes under salt stress. Scientific reports, 5, 1–16.

Singh, A. & Roychoudhury, A. 2020. Role of γ-aminobutyric acid in the mitigation of abiotic stress in plants. Protective chemical agents in the amelioration of plant abiotic stress: biochemical and molecular perspectives, 413-423.

Srivastava, V., Mishra, S., Chowdhary, A. A., Lhamo, S. & Mehrotra, S. 2021. The γ-aminobutyric acid (GABA) towards abiotic stress tolerance. Compatible solutes engineering for crop plants facing climate change, 171-187.

Sun, Y., Oh, D.-H., Duan, L., Ramachandran, P., Ramirez, A., Bartlett, A., Tran, K.-N., Wang, G., Dassanayake, M. & Dinneny, J. R. 2022. Divergence in the ABA gene regulatory network underlies differential growth control. Nature plants, 8, 549–560.

Sundararajan, S., Sivakumar, H. P., Nayeem, S., Rajendran, V., Subiramani, S. & Ramalingam, S. 2021. Influence of exogenous polyamines on somatic embryogenesis and regeneration of fresh and long-term cultures of three elite indica rice cultivars. Cereal Research Communications, 49, 245–253.

Tabur, S., Ozmen, S. & Oney-Birol, S. 2024. Promoter role of putrescine for molecular and biochemical processes under drought stress in barley. Scientific Reports, 14, 19202.

Tamang, B. G., Li, S., Rajasundaram, D., Lamichhane, S. & Fukao, T. 2021. Overlapping and stress-specific transcriptomic and hormonal responses to flooding and drought in soybean. The Plant Journal, 107, 100–117.

Tikhomirova, T. S., Krutovsky, K. V. & Shestibratov, K. A. 2022. Molecular traits for adaptation to drought and salt stress in birch, oak and poplar species. Forests, 14, 7.

Verslues, P. E., Bailey-Serres, J., Brodersen, C., Buckley, T. N., Conti, L., Christmann, A., Dinneny, J. R., Grill, E., Hayes, S. & Heckman, R. W. 2023. Burning questions for a warming and changing world: 15 unknowns in plant abiotic stress. The Plant Cell, 35, 67–108.

Wang, G., Ryu, K. H., Dinneny, A., Carlson, J., Goodstein, D., Lee, J., Oh, D.-H., Oliva, M., Lister, R. & Dinneny, J. R. 2024a. Diversification of gene expression across extremophytes and stress-sensitive species in the Brassicaceae. bioRxiv, 2024.06. 21.599952.

Wang, J., Ding, C., Cui, C., Song, J., Ji, G., Sun, N., Qi, S., Li, J., Xu, Z. & Zhang, H. 2025. Physiological and molecular responses of poplar to salt stress and functional analysis of PagGRXC9 to salt tolerance. Tree Physiology, 45, tpaf039.

Wang, W., Kang, W., Shi, S. & Liu, L. 2024b. Physiological and metabolomic analyses reveal the mechanism by which exogenous spermine improves drought resistance in alfalfa leaves (Medicago sativa L.). Frontiers in Plant Science, 15, 1466493.

Wang, X., Vignjevic, M., Jiang, D., Jacobsen, S. & Wollenweber, B. 2014. Improved tolerance to drought stress after anthesis due to priming before anthesis in wheat (Triticum aestivum L.) var. Vinjett. Journal of experimental botany, 65, 6441–6456.

Wilhelmi, M. D. M. R., Maneejantra, N., Balasubramanian, V. K., Purvine, S. O., Williams, S., Difazio, S., STEWART Jr, C. N., Ahkami, A. H. & Blumwald, E. 2025. Salinity-Induced Photorespiration in Populus Vascular Tissues Facilitate Nitrogen Reallocation. Plant, Cell & Environment, 48, 781–791.

Wuddineh, W., Minocha, R. & Minocha, S. C. 2017. Polyamines in the context of metabolic networks. Polyamines: methods and protocols, 1-23.

Xiong, F., Liao, J., Ma, Y., Wang, Y., Fang, W. & Zhu, X. 2018. The protective effect of exogenous putrescine in the response of tea plants (Camellia sinensis) to salt stress. HortScience, 53, 1640–1646.

Yang, Y. & Guo, Y. 2018. Unraveling salt stress signaling in plants. Journal of integrative plant biology, 60, 796–804.

Yi, L., Wu, M., Yu, F., Song, Q., Zhao, Z., Liao, L. & Tong, J. 2022. Enhanced cadmium phytoremediation capacity of poplar is associated with increased biomass and Cd accumulation under nitrogen deposition conditions. Ecotoxicology and Environmental Safety, 246, 114154.

Yuan, Y., Zhong, M., Shu, S., Du, N., He, L., Yuan, L., Sun, J. & Guo, S. 2015. Effects of exogenous putrescine on leaf anatomy and carbohydrate metabolism in cucumber (Cucumis sativus L.) under salt stress. Journal of Plant Growth Regulation, 34, 451–464.

Zafar, S., Perveen, S., Kamran Khan, M., Shaheen, M. R., Hussain, R., Sarwar, N., Rashid, S., Nafees, M., Farid, G. & Alamri, S. 2022. Effect of zinc nanoparticles seed priming and foliar application on the growth and physio-biochemical indices of spinach (Spinacia oleracea L.) under salt stress. PLoS One, 17, e0263194.

Zalesny Jr, R. S., Stange, C. M. & Birr, B. A. 2019. Survival, height growth, and phytoextraction potential of hybrid poplar and Russian Olive (Elaeagnus Angustifolia L.) established on soils varying in salinity in North Dakota, USA. Forests, 10, 672.

Zhang, H., Zhao, Y. & Zhu, J.-K. 2020a. Thriving under stress: how plants balance growth and the stress response. Developmental Cell, 55, 529–543.

Zhang, K., Khan, M. N., Khan, Z., Luo, T., Zhang, B., Bi, J., Hu, L. & Luo, L. 2024. Seed priming with ascorbic acid and spermidine regulated auxin biosynthesis to promote root growth of rice under drought stress. Frontiers in Plant Science, 15, 1482930.

Zhang, X., Cheng, Z., Zhao, K., Yao, W., Sun, X., Jiang, T. & Zhou, B. 2019. Functional characterization of poplar NAC13 gene in salt tolerance. Plant Science, 281, 1–8.

Zhang, X., He, P., Guo, R., Huang, K. & Huang, X. 2023. Effects of salt stress on root morphology, carbon and nitrogen metabolism, and yield of Tartary buckwheat. Scientific Reports, 13, 12483.

Zhang, Y., Yao, Q., Shi, Y., Li, X., Hou, L., Xing, G. & Ahammed, G. J. 2020b. Elevated CO2 improves antioxidant capacity, ion homeostasis, and polyamine metabolism in tomato seedlings under Ca (NO3) 2-induced salt stress. Scientia Horticulturae, 273, 109644.

Zhao, K., Fan, G., Yao, W., Cheng, Z., Zhou, B. & Jiang, T. 2024a. PagMYB73 enhances salt stress tolerance by regulating reactive oxygen species scavenging and osmotic maintenance in poplar. Industrial Crops and Products, 208, 117893.

Zhao, S., Zhang, Q., Liu, M., Zhou, H., Ma, C. & Wang, P. 2021. Regulation of plant responses to salt stress. International Journal of Molecular Sciences, 22, 4609.

Zhao, X., Wang, S., Guo, F. & Xia, P. 2024b. Genome-wide identification of polyamine metabolism and ethylene synthesis genes in Chenopodium quinoa Willd. and their responses to low-temperature stress. BMC genomics, 25, 370.

Zhu, M., Hsu, C.-W., Peralta Ogorek, L. L., Taylor, I. W., La Cavera, S., Oliveira, D. M., Verma, L., Mehra, P., Mijar, M. & Sadanandom, A. 2025. Single-cell transcriptomics reveal how root tissues adapt to soil stress. Nature, 1-9.

